# A genetically encoded reporter reveals coordinated interferon responses in neurons and non-neuronal cells in the brain

**DOI:** 10.64898/2026.01.08.698513

**Authors:** Sarah R. Anderson, Pailin Chiaranunt, Nicholas M. Mroz, Sarah D. Wade, Fathima N. Pitchai, Shinya Okamura, Ashley McDonough, Yi-Je Chen, Ari B. Molofsky, Jonathan R Weinstein, Raul Andino, Anna V. Molofsky

**Author notes:** Correspondence to: Anna V Molofsky MD PhD, University of California San Francisco, 1550 4^th^ Street, San Francisco, CA 94158, USA., @AnnaMolofskyLab. authors contributed equally (Pailin Chiaranunt, Nicholas M. Mroz). National Center for Biomodels, National Institutes of Applied Research, Taipei, Taiwan.

## Abstract

Interferons (IFNs) are canonical antiviral cytokines with emerging homeostatic functions across tissues, including in the brain. However, detecting IFN-responsive cells remains challenging, particularly in the immune-restricted brain environment, limiting our understanding of the full breadth of IFN signaling across diverse cell types. Here we developed a novel mouse reporter that detects IFN responses in the brain and peripheral tissues, including in neurons. This tool, IFN-brite, <u>B</u>right <u>R</u>eporter of Interferon s<u>T</u>imulated gene <u>E</u>xpression, has two copies of the fluorophore mGreenLantern downstream of the native *Isg15* IFN response gene. We observed IFN responses across multiple cell lineages. Deletion of the Type I interferon receptor *Ifnar1* largely abrogated these responses, including to IFNγ, suggesting that Type II signaling can trigger secondary Type I responses. We observed IFN-responsive cells including neurons in a model of ischemic stroke and during experience-dependent neural plasticity. Furthermore, we found that peripheral infection with Sars-CoV2 but not influenza A leads to IFN-responsive neurons in the brain, suggesting differential impacts of these two pathogens in the nervous system. These data define a broadly useful new tool for studying IFN responsive cells and provide evidence that diverse cell types including neurons can respond to IFNs.

## Introduction

Interferons (IFNs) are an ancient yet highly divergent family of cytokines that co-evolved as an antiviral defense system, but also have many functions beyond host defense ^1,2^. IFN signaling stands out among cytokines for the coherence of the downstream signaling it induces across multiple cell types^3^, suggesting that it can coordinate immune responses within tissues. Despite the fact that the upstream ligands are heterogeneous and signal through multiple different receptors, they all converge on hundreds of IFN stimulated genes (ISGs) downstream of JAK-STAT (Janus Kinase-Signal Transducer and Activator of Transcription) signaling^4,5^. Type I IFNs signal through the IFNAR1/2 heterodimer via ligands that include IFN-β, multiple IFN-α genes, IFN-ε, IFN-κ, and IFN-ω^6^. Type III signaling via IFN-λ uses the same downstream signaling pathways but has a unique receptor subunit (IFNLR1) typically restricted to epithelial cells in barrier tissues and select immune cells^7,8^.Finally, Type ll IFN, a.k.a. IFN-γ, signals through IFNGR1/2. All these IFNs can induce programs of ISGs, some of which are cell-type specific and may be specific to Type I/III vs. Type II signaling. Reliably detecting IFN responses across multiple cell types in their spatial context is critical to defining their functional roles within tissues.

Detecting IFN signaling *in situ* is particularly critical in the brain, a tissue that was once seen as immune privileged^5,9^. IFN responses occur across all central nervous system (CNS) cell types, including oligodendrocytes and microglia in white matter aging^10^; astrocytes in models of inflammation, Alzheimer’s disease, and multiple sclerosis^11,12^; and microglia during developmental remodeling^13^ and models of glioblastoma, stroke, aging, and Alzheimer’s Disease (AD)^13–17^. IFN-responsive microglia are one of the most conserved microglial subsets in human AD datasets^18^. Furthermore, early life viral infection is a strong risk factor for later neurodevelopmental and psychiatric diagnoses ^19,20^, and IFN signaling is dysregulated in a variety of neurodevelopmental disorders (NDDs) including in Type I interferonopathies^1,21–23^, and Down syndrome^24–27^, suggesting key roles of IFN-signaling in brain development.

Defining which cells are IFN-responsive in these and many other biological contexts could yield fundamental insight into the breadth and temporal evolution of these tissue responses. This is particularly true for neurons, given emerging evidence that they undergo neuromodulation by multiple cytokines in both homeostatic and pathological settings, influencing cognition and behavior^28^. For example, IFN-γ promotes sociability by directly signaling to brain inhibitory neurons^29,30^, whereas Type I IFN signaling to both neurons and microglia exacerbate amyloid plaque pathology in an Alzheimer’s Disease (AD) model^31,32^. These findings raise key questions about whether and how neurons respond to IFN and how cellular responses to IFN modulate function. Existing tools to detect IFN responses ^33–35^ reveal the potential of this approach to identify IFN-responsive cell types and suggest the need for a high-sensitivity system that can detect coordinated IFN responses across multiple tissues, including in the brain and other settings where homeostatic and low level IFN responses are prevalent.

Here we report a novel, highly sensitive reporter of IFN responses that can detect IFN signaling across multiple cell types, including in neurons. This *Isg15* knock-in mouse model, IFN-brite, (<u>B</u>right <u>R</u>eporter of Interferon s<u>T</u>imulated gene <u>E</u>xpression) enables *in vivo* and *in vitro* detection of IFN responses and is highly selective for Type I signaling. IFN-brite detected neuronal IFN responses after exogenous cytokines and in a variety of perturbations, including developmental plasticity, ischemic stroke, influenza A infection, and SARS-CoV-2 infection. Notably, IFN-responsive neurons are detected in the brain after SARS-CoV-2 infection but not influenza, revealing a biological use case for this tool. Altogether, our data reveal highly coordinated Type I IFN signaling across multiple cell types within a tissue, including neurons, both after cytokine delivery and in pathophysiological settings.

## Results

### Design and validation of IFN-response reporter, IFN-brite

Using *in silico* analyses we identified *Isg15* as an ISG that met the following criteria: 1) broadly expressed across all known cell types in response to IFN signaling, 2) robustly induced, and 3) not tonically expressed. *Isg15* is a canonical ISG expressed in multiple cell types and based on its genetic sequence and regulatory regions is considered to be preferentially responsive to IFN-I/III relative to IFN-II ^36–38^ (**Fig. 1A**). As a ubiquitin-like protein^39^ its functional role is “ISG-ylation” (a biochemical process similar to ubiquitination that modifies and regulates other ISGs)^40–42^. To generate the knock-in mouse model IFN-brite (<u>B</u>right <u>R</u>eporter of Interferon s<u>T</u>imulated genes), we used CRISPR/Cas9 to insert two copies^43^ of mGreenLantern (mGL), a bright monomeric mutant GFP with a maturation half-time of 13.5 minutes^44^ immediately prior to the stop codon of *Isg15*, to report twofold the fluorescent protein along with the native murine gene (**Fig. 1B**).

**Figure 1:**
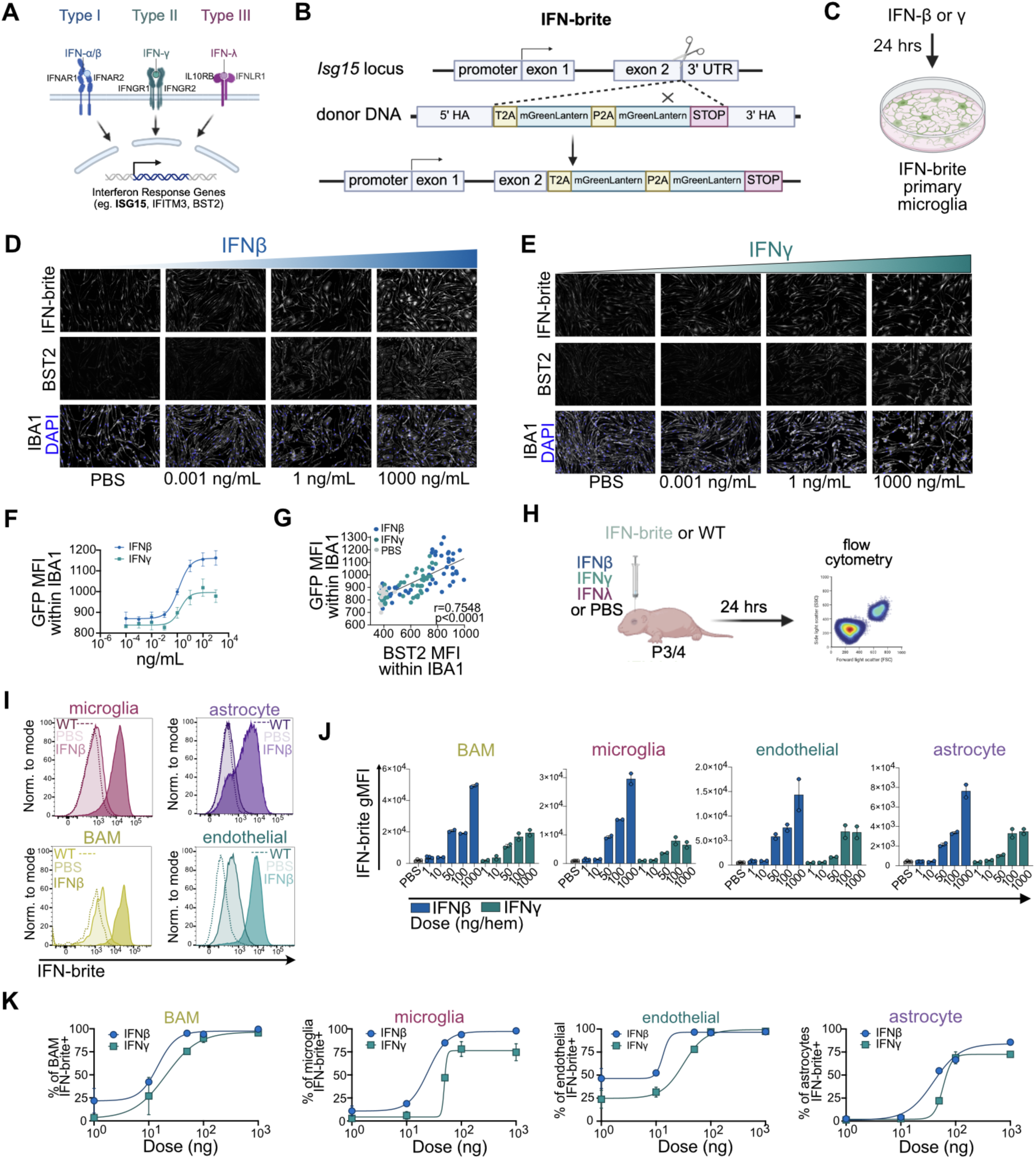
Design and validation of IFN-brite, a high sensitivity IFN-response reporter. A) Schematic of Type I-III IFN signaling pathways that can drive expression of interferon stimulated genes (ISGs), showing select ISGs including *Isg15*. B) Design and generation of IFN-brite reporter. Two copies of mGreen lantern were inserted into the endogenous *Isg15* gene locus. C) *In vitro* assay using mouse primary microglia to detect IFN responses. Cytokines were added for 24 hours prior to fixation, antibody amplification, and imaging. D) Representative images of *in vitro* microglial responses to IFN-β, labelled for IFN-brite (anti-GFP antibody), ISG BST2 and the microglial marker protein IBA1. E) Representative images of *in vitro* microglial responses to IFN-γ using the same imaging panel as in D. F) Best-fit curve of IFN-brite expression in microglia after 24 hours of exposure to IFN-β and IFN-γ at indicated doses by mean fluorescence intensity (MFI) of GFP within IBA1+ mask. n=6 wells, 2 independent experiments per group. Four parameter logistic nonlinear regression, variable slope. IFN-β: EC50=1.295 [0.5917-3.111], hill slope= 0.9625, R^2^=0.8326. IFN-γ: hill slope=1.2, R^2^=0.5353. G) Correlation of IFN-brite expression (anti-GFP) in microglia (IBA1) with BST2. Both quantified as MFI within an IBA1+ mask 24 hours post application. Pearson’s correlation r(106)=0.7548, p<0.0001. Simple linear regression (line): b= 0.5012, F(1,106)= 140.3, p<0.0001, R^2^=0.5697. H) Schematic of *in vivo* dosing strategy to measure IFN-brite responses in CNS cells after bilateral intracerebroventricular (i.c.v.) delivery of interferons. I) Representative histograms showing IFN-brite fluorescence by flow cytometry in the indicated cell types 24 hours after i.c.v. administration. WT, non-transgenic animal after 50ng IFN-β (dashed line); PBS, IFN-brite reporter after vehicle (light shade); IFN-brite reporter after 50ng of IFN-β (dark shade). J) Quantification of IFN-brite geometric mean fluorescence intensity (gMFI) to varying doses of indicated cytokine per cell type. n=2-4 per group. K) Best fit curve of proportion of positive cells at indicated doses for both microglia, astrocytes, BAM, and endothelial cells in response to IFN-β or IFN-γ. N≥2 per dose. Four parameter logistic nonlinear regression, variable slope. BAMs IFN-β:hill slope= 2.581, R^2^=0.962. BAMs IFN-γ: hill slope=1.469, R^2^=0.9673. Microglia IFN-β: hill slope= 2.497, R^2^=0.9947. Microglia IFN-γ: hill slope= 18.84, R^2^=0.9819. Endothelial IFN-β: hill slope= 6.830, R^2^=0.9514. Endothelial IFN-γ: hill slope=2.201, R^2^=0.9847. Astrocyte IFN-β: hill slope= 1.797, R^2^=0.9914. Astrocyte IFN-γ: hill slope= 5.444, R^2^=0.9936. Abbreviations: ISG, interferon-stimulated gene. P, postnatal. i.c.v, intracerebroventricular. BAM, border associated macrophage. gMFI, geometric mean fluorescence intensity. Hem, hemisphere. WT, wild-type.

After validation of founders with correct insertion and germline transmission (**Fig S1A-D** and **Table 1**), we tested sensitivity and specificity to exogenous IFNs in primary microglial cultures incubated 24 hours in exogenous IFN-β, IFN-γ, or IFN-λ2 from heterozygous IFN-brite animals (*Isg15^IFN-brite/+^)* (**Fig. 1C**). We observed robust dose-dependent induction of IFN-brite signal within 24 hours of IFN-β and IFN-γ, although the maximal response to IFN-γ was lower (**Fig. 1D-F**). This effect strongly correlated with protein levels of the ISG BST2^45^ (**Fig. 1G**, R^2^=0.5697, p<0.0001). IFN-brite signal was detected by 16 hours after IFN-β but not IFN-γ (**Fig. S1E)**, raising the question of whether this delayed response may reflect a secondary Type I response.

To determine whether IFN-brite could detect IFN responses *in vivo* within the CNS, we examined expression by flow cytometry 24 hours after intracerebroventricular (i.c.v.) injection of IFN-β and IFN-γ into P3-4 heterozygous IFN-brite (*Isg15^IFN-brite/+^)* pups (**Fig. 1H**), using a cold protease digestion protocol to capture multiple cell types while limiting microglial activation^46^ (gating strategy in **Fig. S1G**). We observed IFN-brite expression in microglia, border associated macrophages (BAM), endothelial cells, and astrocytes, with substantially brighter signal in response to IFN-β vs IFN-γ (**Fig. 1I-J**, total live cells **S1I),** although all cells had detectable expression at higher doses (**Fig. 1K**, example gating in **S1H).** We did not detect any IFN-brite induction in response to IFN-λ2, either in cultured microglia or after i.c.v. delivery *in vivo* (**Fig. S1J-L**), although we did not assess whether receptor protein was detected in these tissues. We did not observe differences in baseline levels of *Isg15* and several other ISGs in IFN-brite animals relative to controls (**Fig. S1M,N**), suggesting that endogenous gene expression is not altered by the insertion. In summary, IFN-brite reports IFN exposure both *in vitro* and *in vivo* and is preferentially sensitive to Type I IFN.

### IFN-brite detects IFN-I responses in neurons *in situ*

We next examined whether IFN-brite could be used for detection *in situ*, including in neurons, which are not amenable to dissociated whole-cell approaches such as flow cytometry. Cytokine injection i.c.v. showed that IFN-β induced robust IFN-brite expression throughout the brain parenchyma, whereas IFN-γ induced weaker and more localized responses around the ventricle and at the injection site at equivalent doses (**Fig. 2A-B**; 50ng/0.5µl per hemisphere). Co-labeling with NeuN revealed IFN-responsive neurons across several brain regions including the anterior cingulate cortex, motor cortex, somatosensory cortex, lateral septum, and caudate putamen (**Fig. 2C**). We found substantially more neuronal IFN-brite after IFN-β than IFN-γ even in regions with the highest amount of neuronal signal (**Fig. 2D**). Therefore, IFN-brite can detect neuronal responses to IFN.

**Figure 2:**
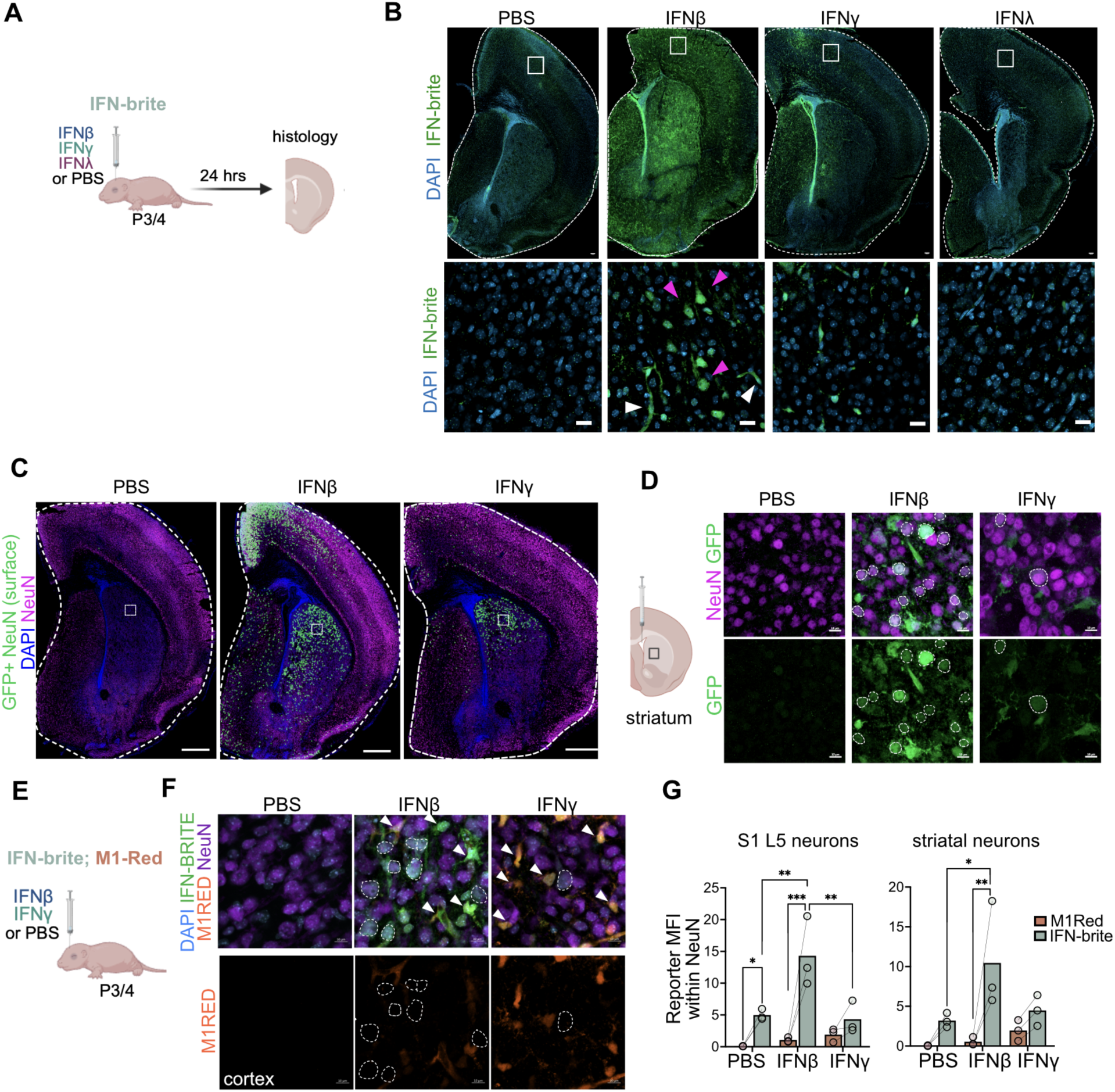
IFN-brite detects IFN-I responses in neurons *in situ*. A) Schematic of *in vivo* dosing strategy to measure IFN-brite responses. B) Representative coronal sections of fixed and antibody stain brain hemisphere at P4-5 showing IFN-brite (GFP) and DAPI (blue). Scale = 100μm top entire coronal section, and 20μm bottom L5 cortex, indicated by boxes. Magenta arrowheads indicate putative neurons. White arrowheads indicate other IFN-responsive cells. C) Representative coronal sections of P4-5 hemispheres 24 hours after i.c.v of indicated ligand. NeuN (purple), masked IFN-brite within NeuN (green), and DAPI (blue). Scale =500μm. White box indicates approximate area of inset in D. D) Representative image of IFN-brite positive neurons in the striatum 24 hours after i.c.v of indicated ligand. Neun (purple), GFP (green). Scale=10μm. White dotted outlines indicate interferon responsive neurons. E) Schematic of *in vivo* dosing strategy of bilateral i.c.v in double reporter, M1Red; IFN-brite with analysis 24 hours later. F) Representative image of S1 cortex 24 hours after i.c.v injection of indicated ligand. GFP (green), NeuN (magenta), DsRed (orange), and DAPI (blue). Scale= 10μm. White dotted outlines indicate interferon-responsive neurons. White arrows indicate interferon responses in non-neuronal cells. G) gMFI quantification of M1Red and IFN-brite signal within NeuN 24 hours after 50ng bilateral i.c.v. of PBS, IFN-β or IFN-γ in S1 cortex (left) and striatum (right). n=3 per condition. (S1 L5 cortex) Two-way RM ANOVA, reporter ***p=0.0008, ligand *p=0.0333, interaction *p=0.0167. Šídák’s multiple comparisons tests: M1Red vs IFN-brite PBS, *p=0.0432, M1Red vs IFN-brite IFN-β, ***p=0.0005; M1Red vs IFN-brite IFN-γ, ns; IFN-brite PBS vs IFN-β **p=0.0023; IFN-γ vs IFN-β, **p=0.0013. (striatum, right) Two-way RM ANOVA, reporter **p=0.0055. Šídák’s multiple comparisons tests: M1Red vs IFN-brite IFN-β, **p=0.0035; IFN-brite PBS vs IFN-β *p=0.0329. Abbreviations: P, postnatal. i.c.v, intracerebroventricular. L, layer. gMFI, geometric mean fluorescence intensity. S1, somatosensory 1 cortex.

To compare IFN-brite to an existing reagent, we crossed it with the Irgm1^DsRed^ (M1Red) reporter^34^ (**Fig. 2E-G, Fig. S2**). We found that both tools could detect responses after IFN administration, and consistent with reports, M1Red preferentially detected IFN-γ (**Fig S2A,B)**. However, IFN-brite was more sensitive overall, detecting many more IFN-responsive neurons in both conditions compared to M1Red (**Fig. 2E-G**). We observed similar patterns in other cell types (**Fig. S3A-J**), whereby IFN-brite was Type I skewed and M1Red was Type II biased, although IFN-brite was 5–35-fold brighter overall (depending on dose), including for IFN-γ, with the exception of ependymal cells of the lateral ventricle (**Fig. S3C-D**). In summary, IFN-brite is a highly sensitive reporter and the only tool to our knowledge that can robustly detect neuronal Type I responses *in vivo*.

### IFN-brite is Type I specific in many cell types at physiological doses

To directly test whether IFN-brite activation was downstream of canonical Type I signaling, we repeated i.c.v. injections in IFN-I receptor knockouts (*Isg15^IFN-brite/+^;Ifnar1^-/-^*). As expected, *Ifnar1* deletion largely abrogated IFN-β-induced IFN-brite expression in all CNS cell types tested, using a half-maximal dose of 50ng (**Fig. 3A-D**). Unexpectedly, *Ifnar1* deletion also reduced IFN-brite expression in response to the same dose of IFN-γ by a similar amount (**Fig. 3B-D**). This could be partly compensated for in some cell types by using a 20-fold higher dose of IFN-γ (1000ng; **Fig S3A**). Neuronal IFN-brite expression in response to IFN-γ showed a similar pattern (**Fig. 3E**), as did ependymal glial cells (**Fig. S3D)**. We also observed loss of ISG15 protein in injected *Ifnar1^-/-^*animals (**Fig.S3C).** Taken together these data indicate that IFN-brite is strongly selective for Type I signaling, and that much of its expression in response to IFN-γ may result from Type I IFN production secondary to IFN-γ signaling.

**Figure 3:**
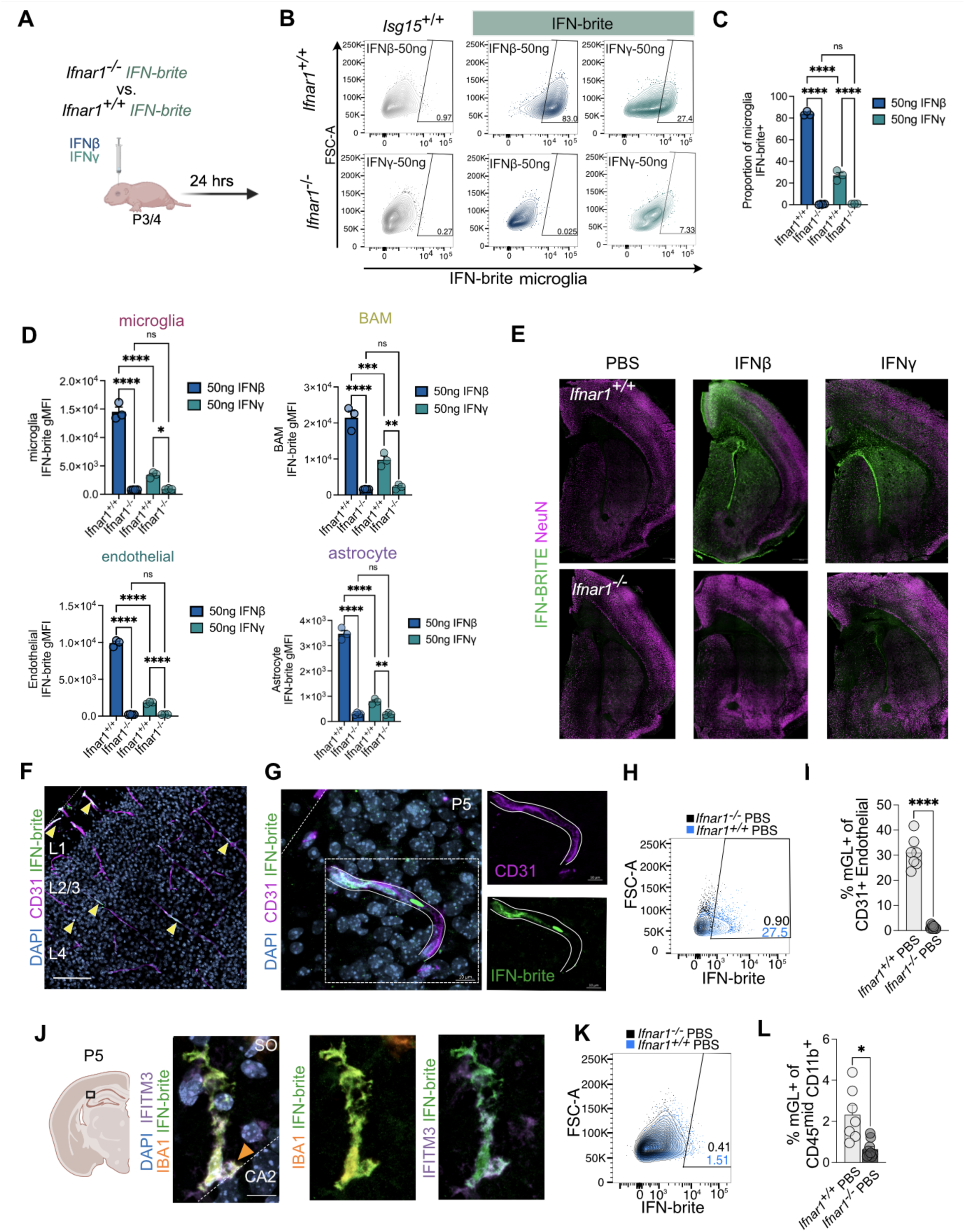
IFN-brite is Type-I specific in many cell types at physiological doses. A) *In vivo* dosing strategy to measure IFN-brite responses in brain cells after i.c.v. delivery of IFN-β in *Ifnar1*^-/-^ and *Ifnar1*^+/+^ *Isg15^Ifn-brite/+^* heterozygous neonates. Analysis 24 hours later. B) Representative contour plots of IFN-brite fluorescence in non-reporter *Isg15^+/+^* (left) or IFN-brite live microglia (right) of *Ifnar1*^+/+^ (top) and *Ifnar1*^-/-^ (bottom) neonates. 24 hours after 50ng of IFN-β (blue) or IFN-γ (green) per hemisphere delivered i.c.v. C) Proportion of IFN-brite^+^ microglia in *Ifnar1^+/+^* and *Ifnar1^-/-^* pups 24 hours after i.c.v IFN-β (blue) or IFN-γ (green) per hemisphere. n=3-4 per group. Two-way ANOVA. Ligand ****p<0.0001, genotype ****p<0.0001. Interaction****p<0.0001. Šídák’s multiple comparisons tests ****p<0.0001. D) IFN-brite intensity (gMFI) of indicated cell types in *Ifnar1^+/+^* and *Ifnar1^-/-^* 24 hours after 50ng per hemisphere dose of IFN-β or IFN-γ. n=3-4 per group. Two-way ANOVA for each. **Microglia** ligand ****p<0.0001, genotype ****p<0.0001. Interaction ****p<0.0001. Šídák’s multiple comparisons tests ****, p<0.0001, *p=0.0129. **BAM** ligand ***p=0.0004, genotype ****p<0.0001. Interaction ***p=0.0002. Šídák’s multiple comparisons tests ****p<0.0001, ***p=0.0001. **p=0.004. **Endothelial** ligand ****p<0.0001, genotype ****p<0.0001. Interaction ****p<0.0001. Šídák’s multiple comparisons tests ****p<0.0001. **Astrocyte** ****p<0.0001, genotype ****p<0.0001. Interaction ****p<0.0001. Šídák’s multiple comparisons tests ****, p<0.0001, **p=0.0065. E) Low power images of *Ifnar1^+/+^* (top) and *Ifnar1^-/-^* (bottom) hemispheres 24 hours after PBS (left) or 50ng IFN-β (right) or 50ng IFN-γ i.c.v. IFN-brite (green) and NeuN (magenta). Scale=500um. F) Representative images of P5 IFN-brite somatosensory cortex (PBS i.c.v) stained with CD31 (magenta) and IFN-brite (green), and DAPI (blue). Scale=100μm. Yellow arrows indicate GFP+ blood vessels. G) High magnification of L1-3 S1 somatosensory cortex in P5 IFN-brite animal (PBS i.c.v). CD31, (magenta), IFN-brite (green) DAPI (blue) with inset (right) outlining walls of blood vessels in individual channels. Scale=10μm. H) Contour plot and gating of IFN-brite^+^ endothelial cells of P4-5 *Ifnar1^+/+^* (blue) and *Ifnar1^-/-^* (black) neonates 24 hours after vehicle, PBS i.c.v. I) Flow cytometry quantification of IFN-brite^+^ endothelial cells in P4-5 *Ifnar1^+/+^* and *Ifnar1^-/-^* neonates. n=7-9 per group. Unpaired two-tailed Welch’s t-test, ****p<0.0001. J) High magnification of interferon-responsive microglia in the naive P5 hippocampal CA2 of IFN-brite neonate. DAPI (blue), ISG IFITM3 (purple), IBA1 (orange), IFN-brite (green). Scale=10μm. Orange arrow indicates DAPI+ inclusion. K) Contour plot and gating of IFN-brite^+^ microglia of P4-5 *Ifnar1^+/+^* (blue) and *Ifnar1^-/-^* (black) neonates 24 hours after vehicle, PBS i.c.v. L) Flow quantification of IFN-brite^+^ live microglia in P4-5 *Ifnar1^+/+^* and *Ifnar1^-/-^*neonates in vehicle condition. n=7-9 per group. Unpaired two-tailed Welch’s t-test, *p<0.0117. Abbreviations: P, postnatal. i.c.v, intracerebroventricular. L, layer. gMFI, geometric mean fluorescence intensity. CA2, Cornu Ammonis 2. SO, stratum oriens.

Our data also suggest that baseline IFN-responses in the developing brain in endothelial cells and microglia are Type I dependent. While we did not detect IFN-responsive neurons in PBS-injected brains, we found that both endothelial cells and microglia had physiologic IFN-responsive subsets. About 30% of all CD31+ endothelial cells in the developing brain were IFN-responsive (**Fig. 3F-I**, S1 cortex) but were absent in *Ifnar1^-/-^* mice (**Fig. 3H,I**). Approximately 2% of total microglia were IFN-brite+ in vehicle-treated animals consistent with our prior findings^13^ (**Fig. 3J-L**). These were significantly reduced in *Ifnar1*^-/-^ animals (**Fig. 2K,L**). In contrast, very low levels of physiologic IFN-brite expression in astrocytes and BAMS were not *Ifnar1* dependent (**Fig. S3B**). In summary, IFN-brite can reveal both homeostatic and IFN-induced responses, and suggests that *in vivo*, responses to IFN-γ may in some cases reflect cross-talk between IFN-I and IFN-II responses.

### IFN-responsive microglia and neurons during development and after ischemic stroke

We previously adapted a model of experience-dependent circuit remodeling that induces a distinct population of IFN-responsive microglia in layer 5 of somatosensory cortex ^13^. The model involves partially lesioning mouse whiskers in early life, which induces rearrangement of the corresponding somatosensory cortex to match the new topographic map (**Fig. 4A**), and leads to expansion of IFN responsive microglia as detected by single-cell RNA sequencing and immunostaining for the ISG, IFITM3 ^13^. To test whether IFN-brite could detect these remodeling-associated microglia we performed whisker-deprivation on IFN-brite animals and co-labeled with IFITM3. We observed an expansion of IFN-brite+ microglia, most of which were also positive for IFITM3 protein (**Fig. 4B-E**). We did not observe any IFITM3+ cells that were not IFN-brite+. In contrast, another IFN reporter mouse (Mx1^GFP^) detected only 25% of IFITM3+ cells^13^. This supported our previous findings and suggested that IFN responses could be detected in response to physiologic perturbations that induce circuit remodeling in the developing CNS.

**Figure 4:**
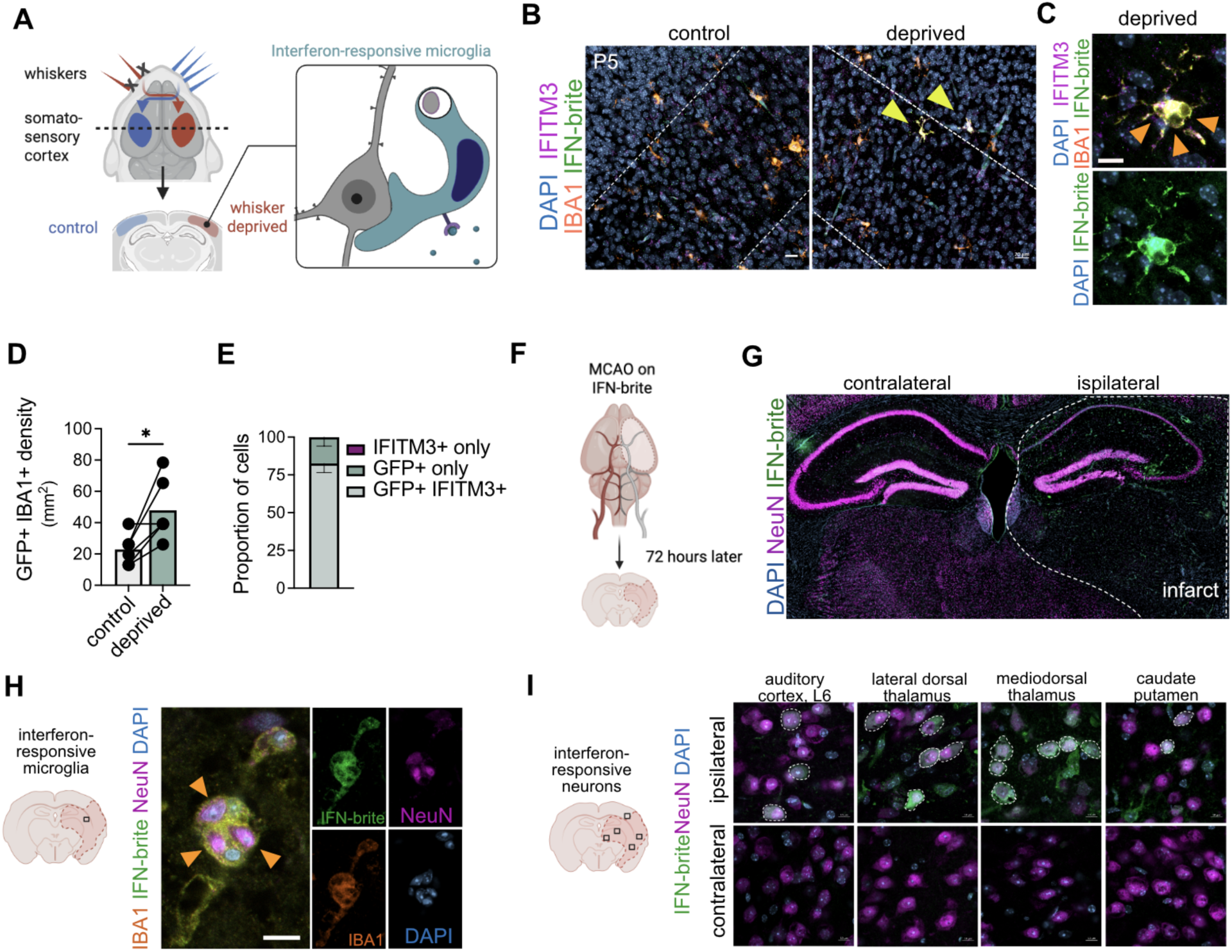
IFN-responsive microglia and neurons during development and after ischemic stroke. A) Model of induced developmental circuit remodeling: partial whisker deprivation in neonatal mice promotes topographic remapping of contralateral somatosensory cortex and induces IFN-responsive microglia (adapted from ^13^). B) Low power images of Layer 5 control and whisker deprived barrel cortex. Yellow arrowheads: IFN-brite^+^IFITM3^+^ microglia. scale= 40μm (left) C) Inset of IFN-responsive microglia in Layer 5 deprived cortex. Orange arrowheads indicate phagosomal compartments. Scale=10μm. D) Quantification of IFN responsive microglia (GFP^+^ IBA1^+^) per mm^2^ in the control and corresponding deprived barrel cortex. n=6. Paired two-tailed t-test *p=0.0218. E) Proportion of IFN responsive microglia single or double positive for IFN-brite (GFP) and existing marker IFITM3. n=6 F) Schematic of transient middle cerebral artery occlusion (tMCAO) on IFN-brite adult mice and analysis 72 hours after reperfusion. G) Representative coronal low power view of hippocampus and thalamus after unilateral tMCAO. IFN-brite (green), NeuN (magenta), and DAPI (blue). Dotted line outlines the infarct region. Scale= 100µm H) Representative high-power image of an IFN responsive microglia after MCAO in dorsal striatum with several phagocytic compartments. Single Z plane (left) and 5um max projection (right). IBA1 (orange), IFN-brite (green), NeuN (magenta), and DAPI (blue). Orange arrowheads indicating engulfed neurons within microglia. Scale=10µm. I) Representative high-power image of interferon responsive neurons across indicated brain regions in ipsilateral (top) and contralateral (bottom). IFN-brite (green), NeuN (magenta), and DAPI (blue). Orange arrowheads indicating engulfed neurons within microglia. Scale=10µm Abbreviations: P, postnatal. L, layer. MCAO, middle cerebral artery occlusion.

We next examined whether IFN-brite can reveal IFN responses after CNS injury. IFN-responsive microglia have been observed in sequencing datasets after ischemic stroke^14,47^, reanalyzed in^13^. IFN-responsive microglia are also induced by ischemic preconditioning and can have protective effects against stroke ^48,49^. To more broadly profile IFN-responsive cells in ischemic stroke we used the transient middle cerebral artery occlusion (tMCAO) model and analyzed 72 hours later (**Fig. 4F**). We found broadly increased IFN-brite signal in the infarcted hemisphere compared to the contralateral hemisphere (**Fig. 4G**). As predicted from the literature, we observed expansion of IFN-responsive microglia particularly in the striatum (**Fig. 4H**). Unexpectedly, we also found many IFN-responsive neurons throughout both the infarct and border-regions on the ipsilateral and not contralateral hemisphere (**Fig. 4I**). Together, these findings suggest that a coordinated IFN-responsive state emerges across multiple cell types, including neurons, after ischemic stroke.

### Interferon responsive neurons in the brain after respiratory viral infection

IFNs are critical for antiviral responses, including after respiratory viral infection^6^. Therefore, we next examined whether IFN-brite could be used to characterize IFN-responsive cell types during two different respiratory viral infections: influenza A virus (IAV) and SARS-CoV-2. First, we infected adult IFN-brite mice with sublethal doses of Influenza A/PR/8/34 (H1N1) (IAV PR8; 50 pfu) (**Fig. 5A**) and collected lungs at 7 days post-infection (dpi) for imaging and flow cytometry. Infected IFN-brite mice did not display significant differences in sickness severity or weight loss compared to infected non-reporter counterparts (**Fig. S4A**).

**Figure 5:**
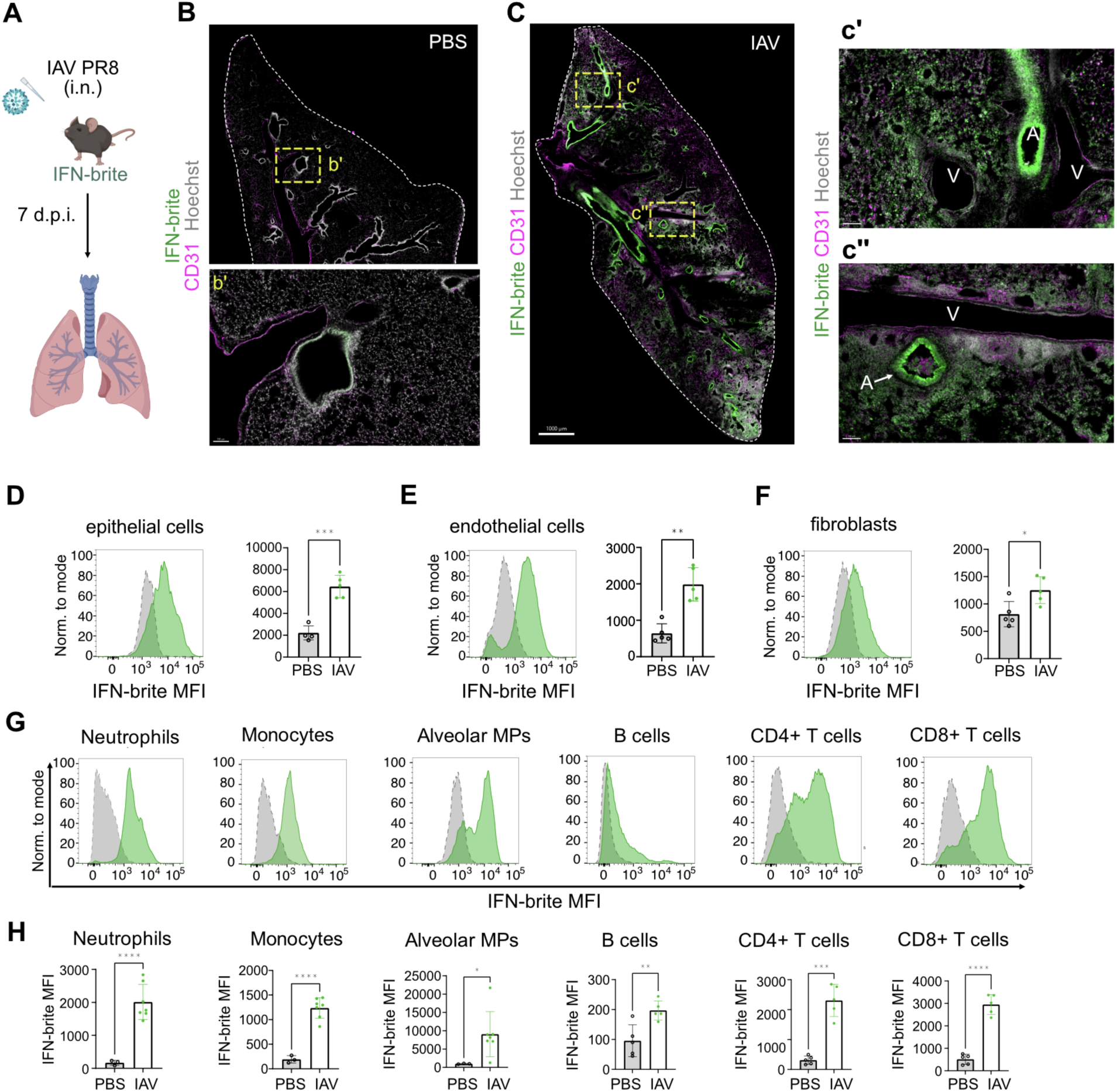
Coordinated IFN responses in lung stromal and hematopoietic cells after Influenza A infection. A) Schematic of influenza A infection paradigm. B) Representative low power image of non-infected lung (PBS vehicle only) in IFN-brite animals. IFN-brite (green), CD31 (magenta), Hoescht (gray). Scale=1mm. Inset shows rare IFN-brite signal at an airway. Scale=100μm. C) Representative low power image and insets of IAV PR8 infected lung 7 days post infection (DPI). IFN-brite (green), CD31 (magenta), Hoescht (gray). Scale=1mm. Insets show IFN-brite airways (A) and parenchyma. V=vessels. D) Histogram and quantification of IFN-brite intensity (MFI) in epithelial cells in vehicle and IAV infected lungs. Data points represent mice from 2 independent experiments; unpaired t-test with Welch’s correction. E) Histogram and quantification of IFN-brite intensity (MFI) in endothelial cells in vehicle and IAV infected lung F) Histogram and quantification of IFN-brite intensity (MFI) in fibroblasts in vehicle and IAV infected lung G) Representative histograms of IFN-brite expression in vehicle (gray) and IAV infected (green) lung immune cell subsets. See Fig. S4 for gating strategy. H) Flow cytometric quantification of IFN-brite in lung immune cell subsets, data points represent mice from 2 independent experiments; unpaired t-test with Welch’s correction. Abbreviations: IAV, influenza A. PR8, Puerto Rico/8/1934 (H1N1). i.n., intranasal. d.p.i, days post infection. MFI, mean fluorescence intensity. A, airways. V, vessel.

We observed robust IFN-brite induction after IAV exposure, including in airway epithelial cells and throughout the lung parenchyma (**Fig. 5B-C**). We quantified IFN-brite induction using flow cytometry, dissociating via enzymatic digestion to collect both immune and stromal subsets (see methods). We observed robust induction of IFN-brite after IAV infection in epithelial and endothelial cells, and in lung hematopoietic cells, including neutrophils, Ly6C^hi^ monocytes, alveolar macrophages, CD8+ T cells, and CD4+ T cells (**Fig. 5G-H**), and to a lesser extent in fibroblasts at this timepoint (**Fig. 5D-F**; gating strategy in **Fig. S4B,C**). In contrast, Mx1^GFP^ detected IFN-responses only in hematopoietic cells after IAV infection^35^, whereas M1Red also detected response in lung alveolar TypeI/II cells^34^. We did not observe significant increases in IFN-brite in splenic hematopoietic cells, except in circulating neutrophils and Ly6C^hi^ monocytes (not shown), consistent with a prior report that IFN response to IAV is primarily confined to the lung and draining lymph nodes^35^. These data suggest that IFN-brite detects IFN responses across multiple immune and stromal cell subsets.

To examine neuronal responses to viral infection, we compared the brains of IAV infected mice to mice infected with a mouse-adapted strain of SARS-CoV-2 (SARS2-N501Y_MA30_; 10^4^ pfu) (**Fig. 6A**) ^50^. At 5 dpi, SARS-CoV-2-infected mice displayed robust IFN-brite induction in the lungs comparable to IAV infection (**Fig. 6B,C**). However, we also observed IFN-brite expression in neurons in SARS-CoV-2 but not in IAV infected mice. The IFN-brite neurons in SARS-CoV-2 were predominantly olfactory neurons within the mitral cell layer, which receive direct input from the olfactory epithelium (**Fig. 6E-F**). We also observed IFN-brite induction along the brain borders in SARS-CoV-2-infected but not IAV-infected mice (**Fig. 6D**). These data are consistent with evidence across multiple species (humans, hamsters, and mice) of SARS-CoV-2 viral transcripts in the brain, and IFN responses in the olfactory bulb and olfactory epithelium ^50–53^. Overall, these data reveal that IFN-brite can detect IFN responses across cell types, organs, and perturbations.

**Figure 6:**
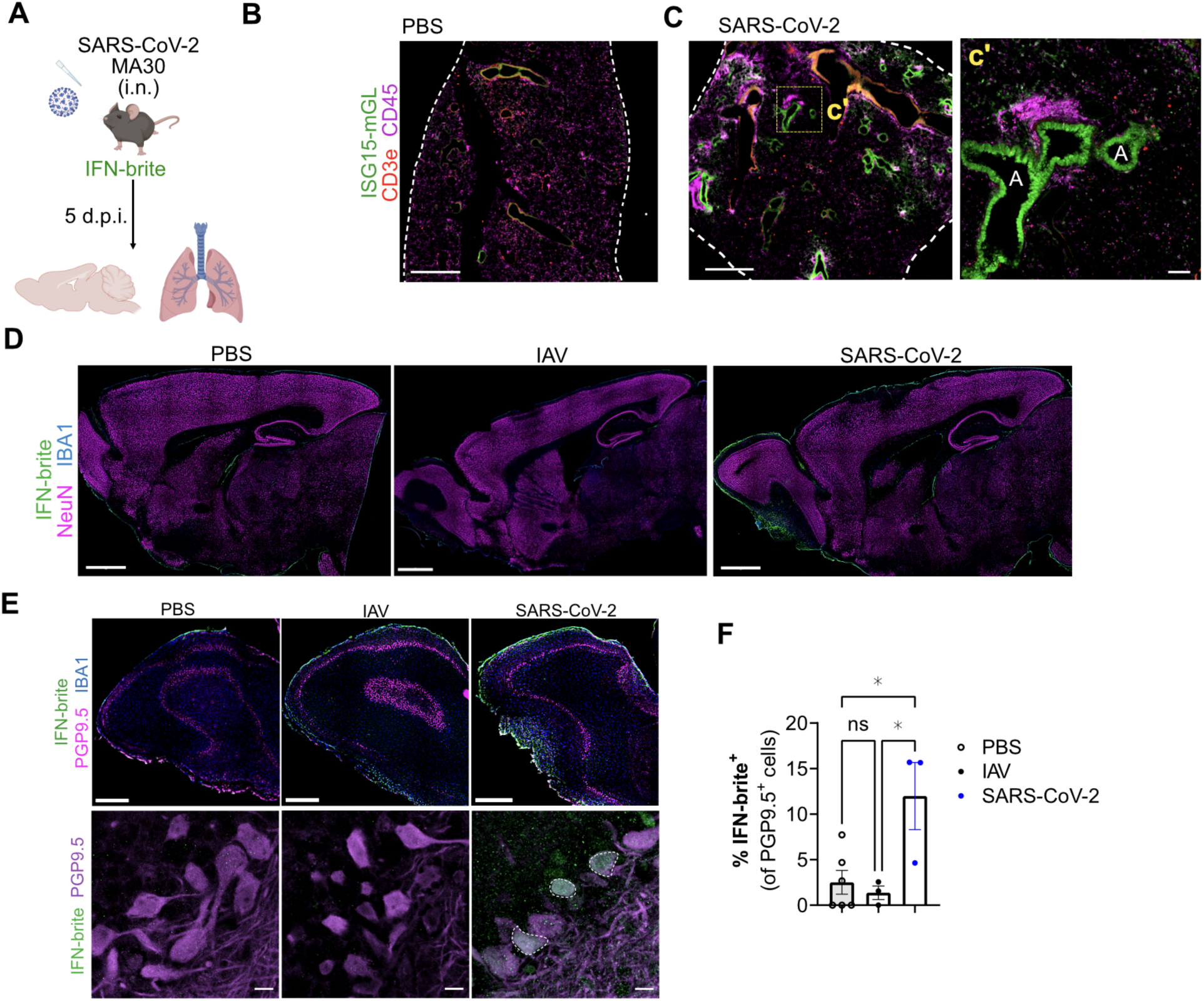
IFN-responsive CNS neurons are detected after SARS-CoV2-2 but not influenza A infection. A) Schematic of SARS-CoV-2 MA30 infection paradigm. B) Representative low power image of non-infected lung (PBS vehicle only) in IFN-brite animals (scale=1mm). IFN-brite (green), CD45 (magenta), CD3e (red). C) Representative low power image and inset of SARS-CoV-2-infected lung 5 days post infection (DPI). IFN-brite (green), CD45 (magenta), CD3e (red). Scale=1mm. Inset shows IFN-brite+ airways, immune cells, and parenchymal cells, with enriched immune infiltration. Scale=100μm. A indicates airway. D) Representative low power widefield images of sagittal brain sections of non-infected (PBS vehicle), IAV-infected (7 DPI), and SARS-CoV-2-infected animals (5 DPI). IFN-brite (green), IBA1 (blue), NeuN (magenta). Scale=1mm E) Representative low power widefield images (top) and high power confocal images (bottom) of the olfactory bulbs of non-infected, IAV-infected, and SARS-CoV-2-infected animals. IFN-brite (green), IBA1 (blue), PGP9.5 (magenta). White dashed outlines indicate IFN-brite+ neurons. Scale=500μm (top), 10μm (bottom). F) Quantification of IFN-brite+ PGP9.5+ neurons in the olfactory bulbs of non-infected, IAV-infected, and SARS-CoV-2-infected animals. One-way ANOVA with Tukey’s test; *p=0.0179 (PBS vs SARS-CoV-2), p=0.0209 (IAV vs SARS-CoV-2); data points represent individual mice. Abbreviations: MA30, mouse adapted SARS-CoV-2. i.n., intranasal. d.p.i, days post infection. A, airways.

## Discussion

Our data suggests that the sensitivity and robustness of IFN-brite allows the determination of coordinated Type I IFN responses in both physiologic and immune activated settings. Furthermore, IFN-brite permits the dissection of cell type- and tissue-specific responses across various contexts including development, injury, infection, and neurodegeneration.

An important finding of our work is that neurons respond to IFN signaling as part of a coordinated IFN response. For example, while ischemic stroke is known to induce IFN-responsive microglial subsets ^13,14,47^, our data suggest that these emerge alongside IFN-responsive neurons, opening new avenues of investigation regarding the relative contributions of these cells in pathology. In addition, we observe IFN-responsive neurons after SARS-CoV-2 but not influenza infection. These two viral pathogens differ in their tropism: influenza A PR8 generally does not cross the blood brain barrier (BBB) or lead to viral proliferation in the brain, whereas SARS-CoV-2 MA30 can^50–53^. Human data suggests that SARS-CoV-2 is more strongly linked to post-viral neurological syndromes and changes in brain volume^54^. Our data suggest that neuronal IFN responses are an additional differentiating factor among these two animal models. Our data are also consistent with a study in AD model mice, which found that both neurons and microglia respond to IFN-I, as loss of *Ifnar1* in these two cell types rescued plaque pathology and synaptic damage via distinct mechanisms^31^. While cytokines signal directly to neurons ^28,55,56^, the extent of these neuromodulatory effects and how they are altered in disease contexts remain largely unknown. IFN-brite may enable assessment of which brain regions or neuronal subtypes are more sensitive to IFN, as well as the mechanistic impact of such signaling.

Our data also indicate that IFN-brite is preferentially tuned to Type I IFN responses. This is consistent with the general consensus in the literature regarding regulation of *Isg15*, which contains two interferon stimulated regulatory elements (ISREs) that drive Type I/III signaling^36,57^ but no evidence of a functional Gamma Activation Site (GAS) element ^58^. It is also consistent with known biological roles of ISG15: intracellularly it restrains IFN-I production^59^ and stabilizes the negative regulator USP18^60^, whereas extracellularly it can act as a cytokine, released in an IFN-β dependent manner, to drive IFN-γ secretion from lymphocytes^61^. As such, Type I specificity might be expected to help avoid a pathogenic positive feedback loop. Although we did not see evidence of Type III responses in our study, this may be due to restricted expression of one of the receptor subunits, IFNL1R. We predict IFN-brite would also be useful for detecting Type III signaling as Type I and III have similar regulation and upregulate a similar set of ISGs including *Isg15*^62,63^.

A secondary implication of the above results is that IFN-γ stimulates secondary Type I signaling. We found an attenuated and temporally delayed IFN-brite response to IFN-γ which was largely eliminated in *Ifnar1^-/-^* mice *in vivo*. Therefore, we hypothesize that IFN-brite responses observed in IFN-γ dominant contexts may be partly due to secondary IFN-⍺/β production driving autocrine or paracrine Type I signaling. The small amount of remaining signal in *Ifnar1^-/-^* mice after IFN-γ could be due to other mechanisms such as IRF1-dependent transcription induced by IFN-γ ^62–64^. This crosstalk has not been reported within the CNS to our knowledge and has interesting implications for IFN regulation and function in CNS health, infection, and disease. It is intriguing that in astrocytes, high doses of IFN-γ could completely overcome *Ifnar1* deficiency, suggesting unique features and regulation of interferon responsive reactive astrocytes (IRRAs). Future studies will further elucidate potential cell type specific mechanisms and biological impacts of this crosstalk within the CNS, as well as determine whether this is broadly true across organs.

Validation of the tool across tissues and by experiments performed at an independent institution (University of Washington) suggest that this reagent should be broadly useful to the field. Our comparisons to an existing reporter suggest that sensitivity or promoter choice may be critical features in enabling detection of IFN-responsive neurons. As with any reporter strain, IFN-brite may have limitations. While we did not observe differences in baseline expression of select ISGs in our study, it is possible that its insertion into the 3’UTR could disrupt regulatory regions or mRNA stability in some contexts. Given that ISG15 partly modulates antiviral responses in response to some pathogens including influenza A^65–67^, it would be prudent to check for any changes in gene expression or phenotype after specific perturbations of interest. Furthermore, the kinetics of expression and half-life of mGreenLantern likely differ compared to ISG15 protein or other ISGs, especially given the prominent negative feedback mechanisms of IFN-I signaling. Therefore, determining temporal dynamics of IFN-brite is an important next step. Development of a nuclear reporter driven by the *Isg15* promoter would allow the isolation of nuclei for important downstream characterization, whereas a conditional tool would enable temporal control of detection. Nonetheless, we expect that this novel resource will enable future studies of IFN cell circuits in the brain and across tissues in physiologic and pathologic settings.

## Author contributions

Conceptualization: S.R.A., A.V.M.

Methodology: S.R.A., P.C., N.M.M., S.D.W., F.N.P, S.O., A.M., Y.C.

Investigation: S.R.A., P.C., N.M.M., S.D.W., F.N.P., S.O., A.M.

Writing - original draft: S.R.A., P.C., N.M.M., A.V.M.

Writing - review & editing: S.R.A., P.C., N.M.M., A.B.M., A.V.M., A.M.

Funding acquisition: A.V.M., A.B.M., J.R.W.

Resources: A.V.M., J.R.W., R.A.

Supervision: A.V.M.

## Acknowledgements

We thank Dr. Richard M. Locksley, Dr. Tien Peng, Dr. Petr Kasparek, and Dr. Zachary Knight for their guidance and contributions. We also thank Junli Zhang and the former Gladstone Transgenic core for valuable technical expertise. We thank Dr. Matthew Spitzer for generous sharing of M1Red mice obtained from the Mutant Mouse Research and Resource Center (MMRRC). We acknowledge and thank the Advanced Light Microscopy at the Weill Innovation Core at UCSF for imaging experiments and the Parnassus Flow Cytometry CoLab and Laboratory for Cell Analysis core at UCSF for flow cytometry experiments.

## Materials and resource availability

IFN-brite mice have been deposited at Jackson Laboratories (JAX, Bar Harbor, ME) under Stock No. 041458. Additional requests for materials or resources to

## Supplementary Figures

**Figure S1:**
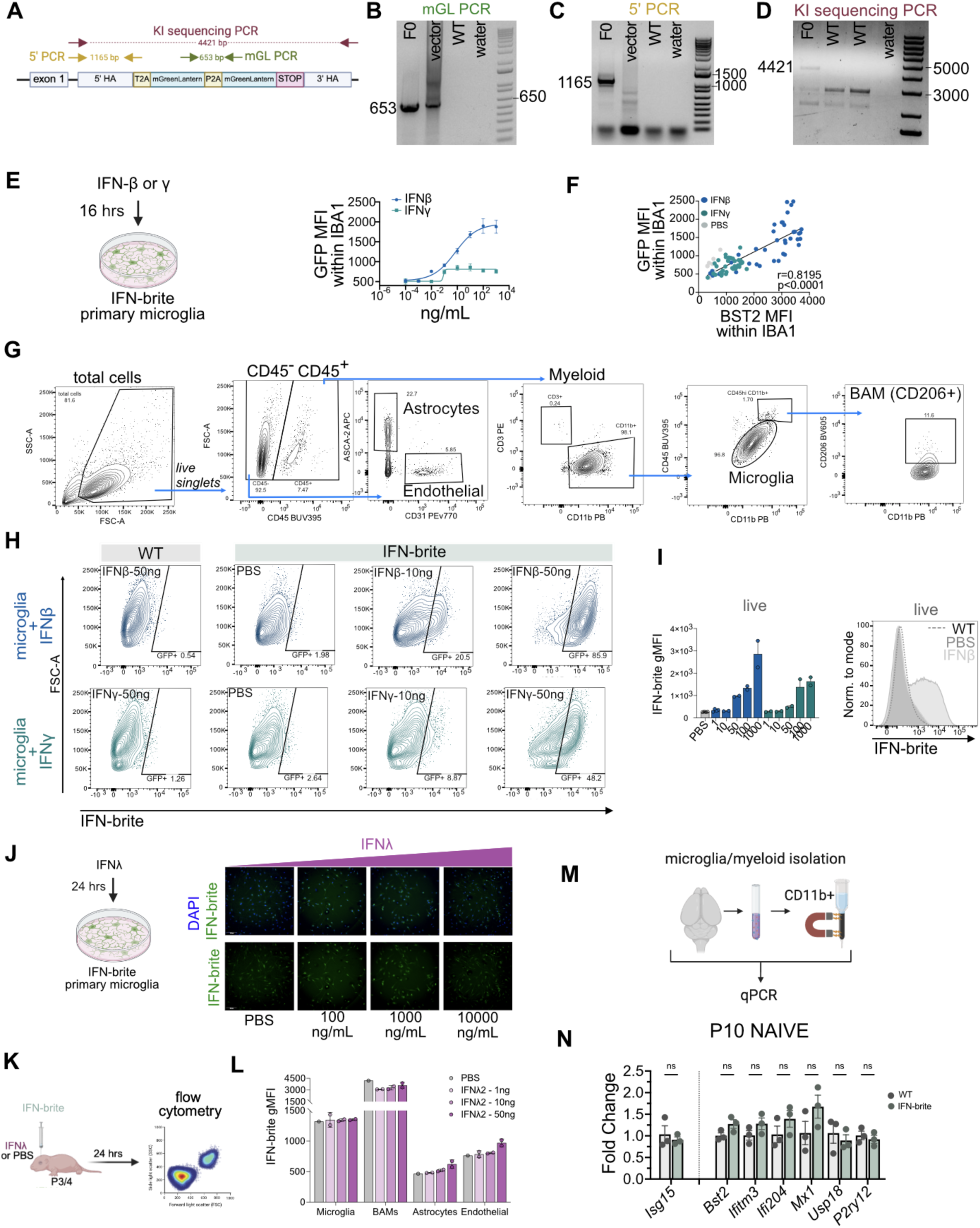
Validation of IFN-brite knock-in and additional controls related to Fig. 1. A) Schematic of IFN-brite with primer sets for validation and genotyping. B) PCR screening of selected F0 pup using mGL PCR primers outlined in A. C) PCR screening of selected F0 pup using 5’ PCR primers outside the homology arm outlined in A. D) PCR screening of selected F0 pup using KI sequencing PCR primers outlined in A. E) Left: experimental design. Right: Best-fit curve of IFN-brite expression in microglia after 16 hours exposure to IFN-β and IFN-γ and by mean fluorescence intensity (MFI) within masked IBA1. n=3-6 wells (0.0001-0.01ng/mL doses n=3), 2 independent experiments per group. Four parameter logistic nonlinear regression, variable slope. IFN-β: EC50=0.7004 [0.1397-3.212], hill slope= 0.5140, R^2^=0.8227. IFN-γ unable to fit. F) Correlation of IFN-brite expression in microglia after protein amplification of mGreenlantern with protein staining of BST2 (a.k.a. tetherin). Both quantified as MFI within an IBA1+ mask 16 hours post application. Pearson’s correlation r(87)=0.8195, p<0.0001. Simple linear regression: b= 0.3795, F(1,87)= 177.9, p<0.0001, R^2^=0.6715. G) Gating strategy for several CNS cell types for all *in vivo* cytokine i.c.v. experiments. H) Contour plots of gating strategy for WT (50ng per hemisphere) and IFN-brite microglia at indicated doses for IFN-β (top) and IFN-γ (bottom). I) The geometric MFI (left) and histogram (right) in total live singlets (Draq^-^) at indicated doses. n≥2 per dose/ligand. J) Left: Schematic of microglia in vitro assay testing IFN-λ2 responses.Right: Representative images of *in vitro* microglial responses to high doses of IFNλ2. DAPI (blue) IFN-brite (green). K) Schematic for IFN-λ2 i.c.v and flow analysis 24 hours later. L) IFN-brite geometric MFI of indicated cell types in vivo at varying doses of IFN-λ2 24 hours after i.c.v injection. n=1 pbs, 2 others. M) Schematic for microglial sorting by magnetic activated cell sorting and qPCR. N) Baseline ISG expression in IFN brite vs littermate control animals. Fold change relative to wildtype littermates of indicated ISGs as well as microglial marker *P2ry12* in naive P10 brain myeloid cells by quantitative reverse transcriptase PCR (normalized to housekeeping gene, *Rps17*), n=3 per group.

**Figure S2:**
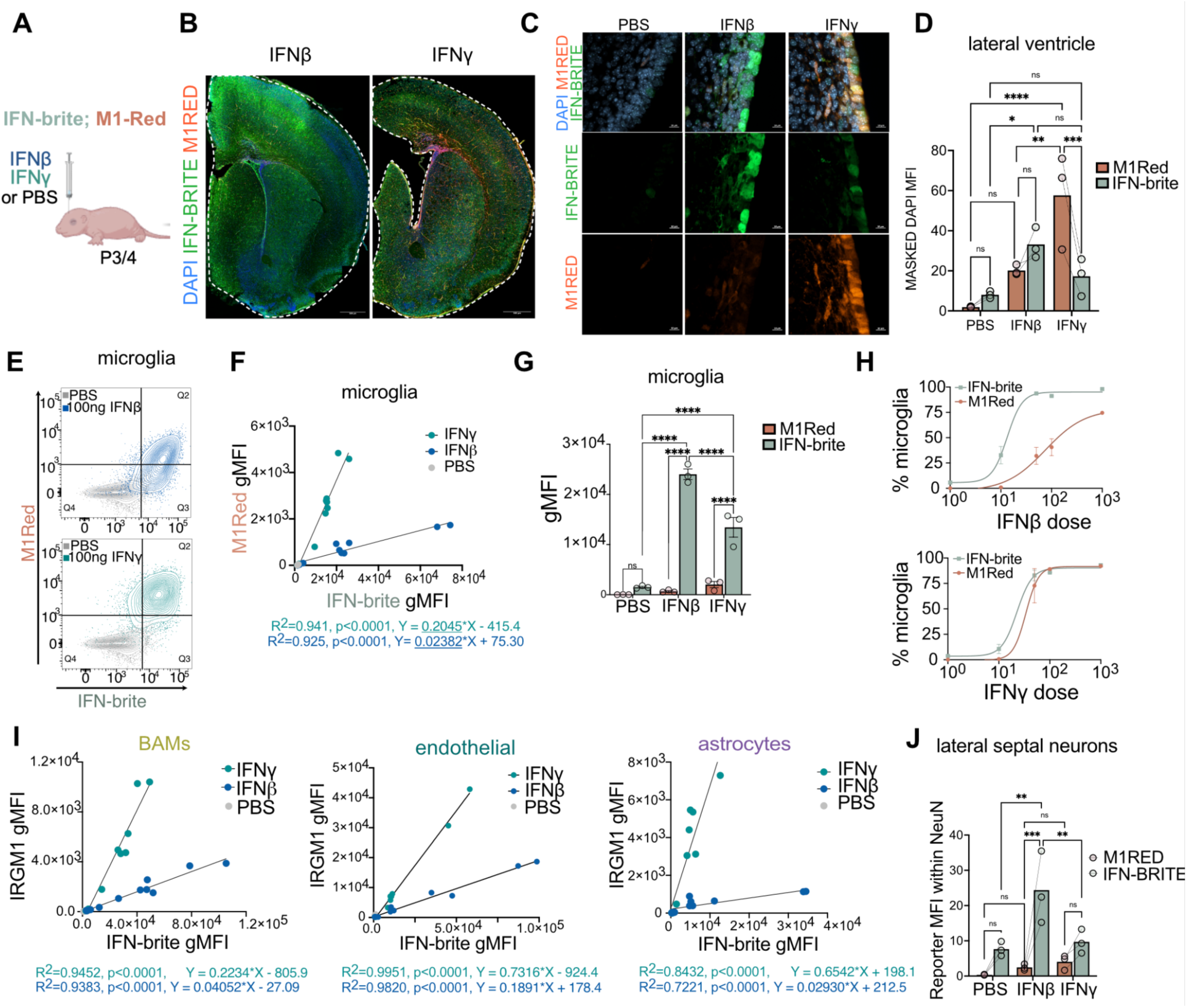
Comparison of IFN-brite to an existing *Irgm1* reporter, related to Figure 2. A) Schematic of *in vivo* dosing strategy in double reporter, M1Red; IFN-brite. B) Low power view of double-reporter 24 hours after i.c.v injection IFN-β or IFN-γ. Scale=500µm. C) Representative low power image of lateral ventricle 24 hours after 50ng IFN-β with inset. IFN-brite (green), IFITM3 (red), DAPI (blue) and IBA1 (magenta). Scale=10µm. D) Quantification of geometric MFI of M1Red and IFN-brite signal within DAPI along the lateral ventricle after 24 hours application of PBS or 50ng per hemisphere of IFN-β or IFN-γ. n=3 per condition. Two-way RM ANOVA, ligand *p=0.0171, interaction **p=0.0014. Uncorrected Fisher’s LSD multiple comparisons tests: M1Red vs IFN-brite IFN-γ ***p=0.0005; IFN-brite PBS vs IFN-β *p=0.0162; M1Red PBS vs IFN-γ****p<0.0001, IFN-γ vs IFN-β **p=0.0013. E) Representative contour plots of M1Red and IFN-brite fluorescence in live microglia in double reporter mice 24 hours after i.c.v. delivery of PBS (gray) or 100ng IFN-β (blue, top) or IFN-γ (green, bottom). F) Scatterplot of gMFI of M1Red and IFN-brite at various doses of PBS (gray), IFN-β (blue), or IFN-γ (green). n= 11 IFN-β, n=12 IFN-γ, n=4 PBS. Simple linear regression, IFN-β: b= 0.0238, F(1,9)= 110.5, p<0.0001, R^2^=0.9247 and IFN-γ: b= 0.2045, F(1,10)= 160.4, p<0.0001 R^2^=0.941. G) Quantification of gMFI of M1Red and IFN-brite of microglia 24 hours after PBS or 50ng per hemisphere of IFN-β or IFN-γ i.c.v. n=3 per condition. Two-way RM ANOVA, reporter ****p<0.0001, ligand ***p=0.0002, interaction ****p<0.0001. Tukey’s multiple comparisons tests: M1Red vs IFN-brite IFN-β, ****p<0.0001; M1Red vs IFN-brite IFN-γ, ****p<0.0001; IFN-brite PBS vs IFN-β; PBS vs IFN-γ, IFN-γ vs IFN-β, all ****p<0.0001. H) Best fit curve of proportion of positive cells at indicated doses for microglia in response to IFN-β or IFN-γ. N≥2 per dose. Four parameter logistic nonlinear regression, variable slope. Microglia IFN-brite IFN-β: hillslope=2.917, R^2^=0.9868. Microglia M1Red IFN-β: hillslope=1.106, R^2^=0.9187. Microglia IFN-brite IFN-γ: hill slope=2.912, R^2^=0.9848. Microglia M1Red IFN-γ: hill slope=3.482, R^2^=0.9129. I) Scatterplot of gMFI of M1Red and IFN-brite at various doses of PBS (gray), IFN-β (blue), or IFN-γ (green) in indicated cell types. n= 11 IFN-β, n=12 IFN-γ, n=4 PBS. Simple linear regression reported below the graphs. J) Quantification of neurons in double reporter animals after i.c.v. cytokine delivery in lateral septum. n=3 per condition. Two-way RM ANOVA. reporter ***p=0.0009, ligand *p=0.0449, interaction *p=0.0245. Šídák’s multiple comparisons tests: M1Red vs IFN-brite IFN-β, **p=0.0006. IFN-brite PBS vs IFN-β **p=0.0023, IFN-β vs IFN-γ **p=0.006.

**Figure S3:**
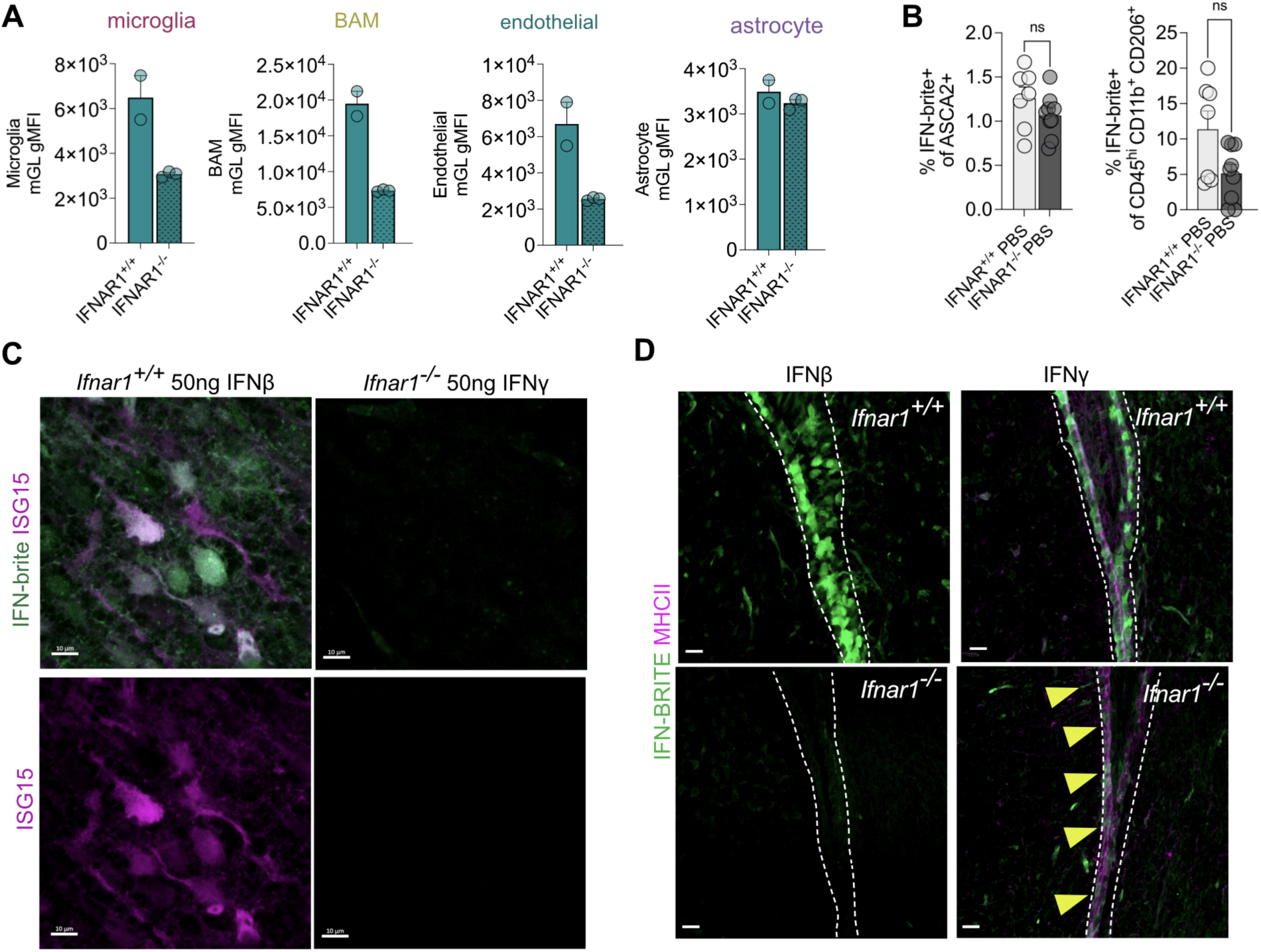
Additional validation of IFN-brite reporter. Related to Figure 3. A) IFN-brite intensity (gMFI) of indicated cell types in *Ifnar1^+/+^* and *Ifnar1^-/-^* 24 hours after 1000 ng per hemisphere dose of IFN-β or IFN-γ. n=2-4 per group. B) Proportion of ASCA2^+^ astrocytes IFN-brite positive (left) and BAM (right) in *Ifnar1^+/+^* and *Ifnar1^-/-^* 24 hours after PBS i.c.v. Unpaired Welch’s t test. C) Representative images of L5 S1 cortex of IFN-brite (green) animals stained with ISG15 antibody (magenta) in Ifnar1^+/+^ 50ng IFN-β (left) or Ifnar1^-/-^ 50ng IFN-γ (right). Scale=10µm. D) Representative low power image of lateral ventricle 24 hours after 50ng IFN-β (left) or 50ng IFN-γ (right) in *Ifnar1^+/+^* (top) and *Ifnar1^-/-^* (bottom) animals. IFN-brite (green) and MHC-II (magenta). Scale=10µm. Dotted lines outline ventricle. Yellow arrows indicate IFN-brite ventricular cells.

**Figure S4:**
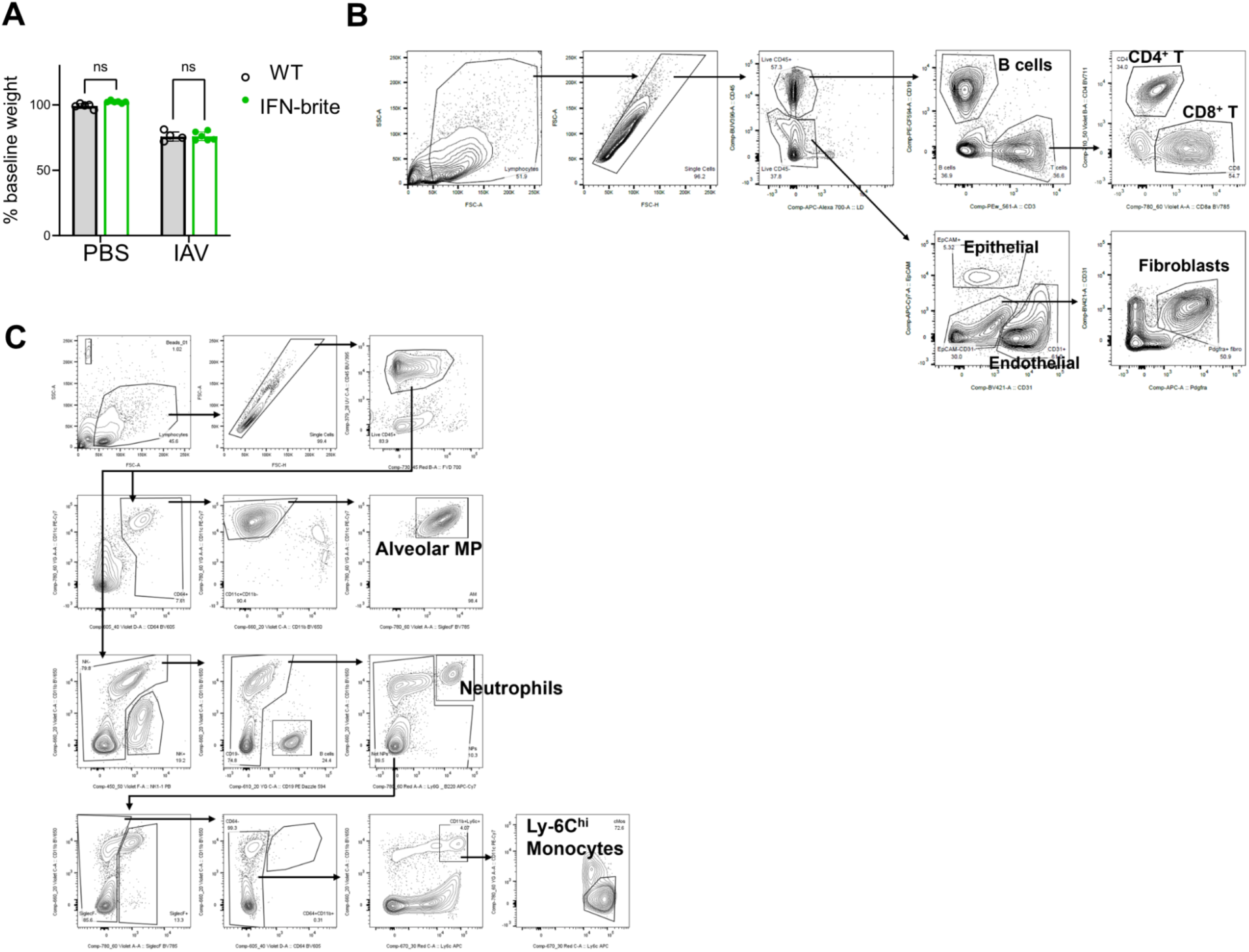
Influenza A infection severity in IFN-brite mice, and gating strategy for lung flow cytometry, related to Fig. 5 A) Body weight (as percent of baseline day 0 weights) of PBS-treated versus IAV-infected WT and IFN-brite mice at 8 dpi. n=5-6 per group, 2 independent experiments, two-way ANOVA with Sidak’s multiple comparisons test. B) Gating strategy for stromal and lymphoid cells analyzed in Fig. 5. C) Gating strategy for myeloid cells analyzed in Fig. 5. (MP = macrophages)

## Brief methods

### Mice

All mouse strains were maintained in the University of California, San Francisco specific pathogen–free animal facility, and all animal protocols were approved by and in accordance with the guidelines established by the Institutional Animal Care and Use Committee and Laboratory Animal Resource Center. All IAV PR8/SARS-CoV-2 infection experiments were performed in a BSL2/BSL2+ facility, respectively. Littermate controls were used for all experiments when feasible, and all mice were backcrossed > 10 generations on a C57BL/6 background unless otherwise indicated. IFN-brite animals were backcrossed for 3 generations prior to experiments to breed out potential indels. All experiments were performed in heterozygous animals and incorporated animals of both sexes in approximately equal numbers. M1Red mice^34^ (#06913-JAX) were obtained from the Mutant Mouse Resource and Research Centers (MMRRC) (submitted by Dr. Alan Sherr and Dr. Lionel Feigenbaum) via collaboration with Dr. Matthew Spitzer (UCSF) and were bred in-house with IFN-brite mice for experiments.

### Generation of IFN-brite animals

The donor DNA was purchased and linearized by Vectorbuilder (Chicago, IL). The ∼4kb donor DNA including homology arms, two copies of mutant mGreenLantern separated by T2A and P2A sites (**Fig. S1A**) was dialyzed using millipore membrane filter (CAT# VSWP04700). CrispR guide, GGAGGGGACCAGTGTGCCTA, was selected based on proximity to the insertion site near the *Isg15* stop codon and purchased from IDT (Newark, New Jersey). crRNA and tracrRNA were prepared in house. RNA and donor DNA was injected along with Cas9 by the Gladstone Transgenic Core. A total of 105 embryos were implanted, thirteen pups were born, and twelve pups survived and were screened (**Fig. S1A-D**). Eight of the twelve had insertions of the donor DNA, five of which were inserted into the correct locus. All five were sequenced and the F0 was selected based on lack of mutations and confirmation of germline transmission (**Table 1**).

### Genotyping primers

Common F: 5’-CTG GTG AGG AAC GAA AGG GG-3’

WT R: 5’-AAA GGG TCT GGA AGG GGC AT-3’

2xmGL R: 5’-TCG ACG TCA CCG CAT GTT-3’

Product sizes of WT= 520 bp and 2xmGL= 284 bp.

### Isolation and culture of primary glia cells

Mixed glia cultures were generated from P2-P4 pups from heterozygous IFN-brite and grown in T75 flasks with DMEM supplemented with heat inactivated 10% FBS and 1% Pen/Strep at 37°C 5% CO2. The media was changed the next day and cultures were grown for another 10-12 days. Microglia were detached by hitting the flasks 10x against the bench, stained with Trypan blue and counted using a Countess 3 Automated Cell Counter (Invitrogen). Then cells were plated in a 96-well plate at a density of 30,000 cells/well in 200ul final volume (3-6 replicate wells per condition). The next day, microglia were treated with vehicle or recombinant cytokine at the indicated dosages, then fixed in 4% paraformaldehyde after either 16 or 24 hours incubation and immunostained.

### Tissue preparation for immunostaining

Mice were anesthetized and perfused transcardially with PBS followed by 4% paraformaldehyde (PFA) fixative for histology only experiments. For some experiments, mice were perfused with PBS only and brains hemisected-one hemisphere used for flow cytometry and one hemisphere dropfixed for 24 hours for immunostaining. Tissues were post-fixed overnight and then transferred to a 20-30% sucrose solution for a minimum of 2 days. Tissues were then embedded in OCT (VWR 25608-930), frozen on dry ice, and stored at −80°C until sectioning. For immunohistochemistry experiments, floating brain sections of 40 µm thickness or lung sections of 200 µm thickness were cut using a Cryostar NX70 cryostat (Thermo).

### Immunohistochemistry, confocal microscopy, and image acquisition

For immunohistochemistry, floating brain sections (30 um, 40 µm, 50µm, or 300 µm in thickness) were incubated in the Staining Solution (0.4% TritonX-100, 5% normal goat or donkey serum, 1X PBS) for one hour at room temperature. Primary antibodies were diluted in Staining Solution, and tissue was incubated in primary solution on a shaker overnight at 4°C or room temperature. Secondary antibodies were diluted in Staining Solution, and tissue was incubated on a shaker for 2 hours at room temperature. Hoechst 33342 (Thermo Fisher) was added at 1:5000 during the last 5 minutes of incubation. Brain sections were mounted on slides and sealed onto coverslips with ProLong Glass without DAPI (Thermo) for high-resolution imaging, SlowFade Glass (Thermo) for 300 µm section immunostaining, and Fluoromount-G (SouthernBiotech) for all other experiments.

For lung immunohistochemistry, floating lung sections (200 µm in thickness) were washed and incubated in permeabilization buffer (DPBS/0.2%Triton X-100/0.3M glycine) for 4-5 hours at room temperature, then blocked in DPBS/0.2% TritonX-100/5% serum (from the same host species as the secondary antibody) at room temperature overnight. After, samples were washed in DPBS/0.2% Tween-20 once and incubated with primary antibodies diluted in DPBS/0.2% Tween-20/5% serum, room temperature until the next day. Next, samples were washed in DPBS/0.2% Tween-20 for 30 min, 3–4 times, then incubated with secondary antibodies diluted in DPBS/0.2%Tween-20/5% serum at room temperature overnight. Samples were washed in PBS for 2hrs, then cleared by incubating in refractive index matching solution (RIMS; 80% Histodenz in 1X PBS, 0.01% sodium azide, 0.1% Tween20) until transparent and then mounted in fresh RIMS solution.

Several GFP antibodies were tested for reactivity to mGreenLantern. Chicken anti-GFP (Aves lab GFP-1020), Guinea pig anti-GFP (Synaptic systems 132 004) and goat anti-GFP (Abcam ab6673) successfully amplified mGreenlantern signal in organs tested while rabbit anti-GFP (Cell signaling technology 2956S) did not.

High resolution images were captured on a Zeiss LSM 800 confocal microscope at 20x, 40x, and 63x objectives. Whole brain or lung confocal images were acquired on a Stellaris 8 confocal microscope (Leica Microsystems) at either 10x (PL APO 10x/0,4 NA) or 20x (PL APO 20x/0.75 NA) of a single Z plane at 1024×1024 with 2x line averaging. *In vitro* imaging was performed on a Molecular Device ImageXpress Confocal HT.ai High Content Microscope at 20X water immersion objective (Apo LWD 20x/0.95NA WI) of a single Z plane. 9 images per well were acquired, analyzed, and averaged. For widefield images, a Leica DMi8 Inverted microscope (Leica microsystems) equipped with a multichannel LED and an ORCA Flash4 sCMOS camera (Hamamatus) at 10x objective (PL APO 10x/0.45) using the LASX software. Stellaris, Widefield, and molecular device confocal was provided by the Advanced Light Microscopy at the Weill innovation core. Within experiments, image settings were consistent across experimental groups. All images were acquired with sequential line scanning, bidirectionally. Pinhole sizes were set to 1 Airy unit for all channels. All images were acquired in 8-bit mode. For lung, slides were imaged on a Stellaris 8 confocal microscope (Leica microsystems) at 10x and 20x objectives. Laser powers and gain were consistent within experiments.

Images were analyzed using ImageJ2, version 2.14.0/1.54f (NIH). 3-4 sections were imaged per animal per brain region and averaged. Mean fluorescence intensity (MFI) within NeuN was determined by creating a threshold mask (Otsu) of positive NeuN signal first and measuring the MFI of either GFP or dsRED signal within each neuron ROI and then averaged per mouse per brain region. MFI within the lateral ventricle was performed by creating a threshold mask (Percentile) on DAPI signal and then measuring the GFP or dsRED MFI within. The density of GFP+ or IFITM3+ IBA1 cells was analyzed from 20x images, 2-3 sections per animal within the barrel cortex and averaged per mouse. Animals of both sexes were used for analysis. For representation of IFN-brite^+^ neurons, Imaris software (Oxford Instruments Andor, Belfast, Ireland) was used to create a surface based on NeuN mean fluorescence intensity and then filtered on IFN-brite mean fluorescence intensity (10-150) with exact settings applied across conditions.

### Stereotaxic injections

All brain injections were performed with a Kopf stereotaxic apparatus (David Kopf, Tujunga, CA) and a microdispensing pump (World Precision Instruments) holding a Hamilton Syringe (model 701 RN, 10 ml) with a beveled glass needle (∼50 mm outer diameter). For intraventricular injections into neonates (P0-2), mice were anesthetized by hypothermia on ice for 3-4 minutes. Bilateral i.c.v. injection of cytokine was performed (from lambda: 1.5 mm AP, +/- 0.8 mm ML, −1.75 mm DV). Afterwards, pups were warmed on a heating pad until full recovery before returning to their home cage. Carrier-free IFN-β was purchased from R&D (Cat#8234-MB-010/CF) and reconstituted and diluted in sterile DPBS. For carrier-free IFN-γ injections, Biolegend (Cat#575304) was used for concentrations below 50ng and R&D (Cat#485-MI-100/CF) was used for higher concentrations (100-1000ng).

### Brain dissociation for flow cytometry

P4-5 mice were deeply anesthetized via hypothermia and transcardially perfused with 5-10 mL of ice cold 1X PBS via the left ventricle. Brains were isolated, hemi-sected, and collected into 2 mL of iMED+ [isolation medium: 1X HBSS (without calcium or magnesium) supplemented with 10 mg/mL phenol red, 15 mM HEPES, and 0.6% glucose]. Hemibrain samples were then homogenized using a “cold protease” protocol optimized to isolate astrocytes, endothelial cells, and hematopoietic cells (including microglia), adapted from ^46^. Samples were digested on ice for 20-25 minutes in 1X DPBS (no calcium or magnesium) supplemented with 5 mg/mL subtilisin A (Protease from *Bacillus licheniformus,* Sigma #P5380) and 20 μg/mL DNase I (Roche #10104159001) with repeated trituration using a P1000 pipette every 5 minutes. Once homogenized, digestion was halted using a solution containing protease inhibitors (“LO-OVO”: 1X HBSS without calcium/magnesium supplemented with 10 mg/mL phenol red, 0.08% glucose, 1 mg/mL ovomucoid (Worthington), 1 mg/mL bovine serum albumin (Sigma), 20 μg/mL DNAse

I). Digested samples were filtered through a 70 μm cell strainer and cells were collected via centrifugation at 300xg for 10 minutes at 4°C. Cell pellets were then resuspended in a 22% Percoll solution (GE Healthcare) with a 1X DPBS layer floated on top, as previously described ^68^. Samples were centrifuged at 900xg for 20 min (acceleration=4, brake=0) at 4°C and the supernatant layers were discarded to remove myelin and debris. Cell pellets were resuspended prior to antibody staining for flow cytometry.

### Lung dissociation for flow cytometry

Single cell suspensions were prepared from lung and spleen tissues as described previously^69–71^. Briefly, mice were euthanized with CO_2_. Immediately after, mice were perfused by flushing the left ventricle with 10 mL 1X DPBS, and whole lungs (left lobe) were excised. Tissues were minced into small pieces (<1 mm). For flow cytometry of immune panels, samples were digested in 1X Hanks’ Balanced Salt Solution (HBSS) with 0.2 mg/mL Liberase Tm (Roche, Cat# 5401127001) and 25 μg/mL DNase 1 (Roche, Cat# 10104159001) for 30 min at 37°C on a shaker. For flow cytometry of stromal cells, samples were digested in PBS with 7.5 U/mL Dispase II (Gibco, ThermoFisher Cat# 17105041), 112 U/mL Collagenase I (Gibco, ThermoFisher Cat# 17100017), and 40 μg/mL DNase 1 (Roche, Cat# 10104159001) for 30 min at 37°C on a shaker. Tissue pieces were then sheared through a 18g needle, followed by filtration through 70μm filters, washed, and subjected to red blood cell lysis (1X Pharm-Lyse lysing solution; BD Biosciences) before final suspension in FACS buffer (1X DPBS, 3% FBS, 0.05% NaN_3_).

### Staining for flow cytometry

For brain samples, single cell suspensions were stained in iMED-(1X HBSS without calcium/magnesium/phenol red supplemented with 15 mM HEPES and 0.6% glucose) containing fluorophore-conjugated antibodies, viability dye (Draq7, 1:1000, Biolegend), and Fc Block (2.4G2, 1:100, BD Biosciences) for 45 min at 4°C. Cells were washed with iMED-, pelleted at 300xg for 5 min at 4°C, and resuspended in iMED-prior to analysis. Data was collected on a FACSAria Fusion (BD).

For lung samples, single cell suspensions were stained in FACS buffer (1X DPBS, 3% FBS, 0.05% NaN_3_) containing fluorophore-conjugated antibodies, viability dye (Draq7, 1:1000, Biolegend), and Fc Block (2.4G2, 1:100, BD Biosciences) for 45 min at 4°C. Cells were washed with FACS buffer, pelleted at 1200rpm for 2 min at 4°C, and resuspended in FACS buffer prior to analysis.

Data was collected on a dual Fortessa X20 (BD) in the Parnassus Flow Cytometry CoLab or FACSAria Fusion (BD) provided by the Laboratory for Cell Analysis core at UCSF. Analysis of flow cytometry data was performed using FlowJo v10 software (BD).

### Neonatal whisker lesion

Postnatal day 2 pups were anesthetized via hypothermia (3-4 minutes on ice). Two incisions were made of rows B and D of the right whisker pad and hair follicles were removed. Silver nitrate was used to cauterize the exposed follicle and prevent regrowth. This was followed by topical application of 2% lidocaine for pain management and recovery on a circulating water pad for at least 15 minutes prior to being returned to their home cage. Animals were collected for perfusion 72 hours later at postnatal day 5 as we have previously done^13^.

### Transient middle cerebral artery occlusion (tMCAO)

The experimental study was designed following National Institutes of Health (NIH) guidelines and with the approval of the University of Washington Institutional Animal Care and Use Committee. IFN-brite animals 21 weeks of age received a transient (60 minute) middle cerebral artery occlusion surgery followed by 72 hours of reperfusion at University of Washington as previously published following the Longa method^48,72^. Animals were anesthetized under isoflurane and occlusion was performed using the transient intraluminal filament method with laser Doppler flowmetry to confirm occlusion (at least 70% reduction in cerebral blood flow) and reperfusion as we have done previously; all mice met our inclusion criteria. Incisions were sutured closed and mice were allowed to recover for 72 hours prior to collection of tissue for analysis.

### Influenza A and SARS-CoV-2 infection

Mice were anesthetized with isoflurane. 50 pfu of Influenza A/PR/8/34 (H1N1) (ATCC, VR-95) or 10^4^ pfu of SARS-CoV-2 (SARS2-N501Y_MA30_) was administered intranasally through each nostril (for a total of 40 μl volume) using a P200 pipette. After administration, mice were held with nostrils upright for 10–15 s to ensure complete inhalation of the virus. Infected mice were monitored daily from the day of infection until the specified time points for tissue collection. All experiments with IAV or SARS-CoV-2 were performed in a biosafety level 2 (BSL2) and BSL2+ laboratory, respectively.

### Quantification and statistical analyses

Graphpad Prism 9.4.1 was used for most statistical analyses. Statistical tests are as described in the text and figure legends.

### Antibodies for flow cytometry

*Antibodies for flow cytometry:*
| **Antigen–**<br>**Fluorophore** | **Clone** | **Dilution** | **Vendor** | **Catalog #** |
| --- | --- | --- | --- | --- |
| CD11b–Pacific Blue | M1/70 | 1:200 | Biolegend | 101224 |
| CD45–BUV395 | 30-F11 | 1:400 | BD Biosciences | 564279 |
| ASCA-2–APC | IH3-18A3 | 1:100 | Milltenyi Biotec | 130-117-386 |
| CD31–PE-vio770 | 390 | 1:100 | Milltenyi Biotec | 130-102-375 |
| CD206–BV605 | C068C2 | 1:200 | Biolegend | 141721 |
| CD3–PE | 17A2 | 1:200 | Biolegend | 100206 |
| CD31–BV421 | 390 | 1:400 | Biolegend | 102424 |
| Sca-1–BV711 | D7 | 1:200 | Biolegend | 108131 |
| CD8α–BV785 | 53-6.7 | 1:200 | Biolegend | 100750 |
| PDGFRα–APC | APA5 | 1:200 | Biolegend | 135908 |
| EpCAM–<br>APC-Fire750 | G8.8 | 1:200 | Biolegend | 118230 |
| CD4–BV711 | RM4-5 | 1:200 | Biolegend | 100557 |
| TCRγδ–<br>PerCP-Cy5.5 | GL3 | 1:200 | Biolegend | 118118 |
| Ly-6C–BV605 | HK1.4 | 1:400 | Biolegend | 128035 |
| NK1.1–BV650 | PK136 | 1:200 | Biolegend | 108736 |
| CD64–AF647 | X54-5/7.1 | 1:200 | BD Biosciences | 558539 |
| Ly-6G–APC-Cy7 | 1A8 | 1:200 | Biolegend | 127624 |
| MHCII–BUV737 | M5/114.15.2 | 1:400 | BD Biosciences | 748845 |
| CD19–<br>PE-Dazzle594 | 6D5 | 1:400 | Biolegend | 115554 |
| CD11c–PE-Cy7 | N418 | 1:200 | Biolegend | 117318 |
| Siglec-F–BV786 | E50-2440 | 1:200 | BD Biosciences | 740956 |

### Antibodies for immunofluorescence

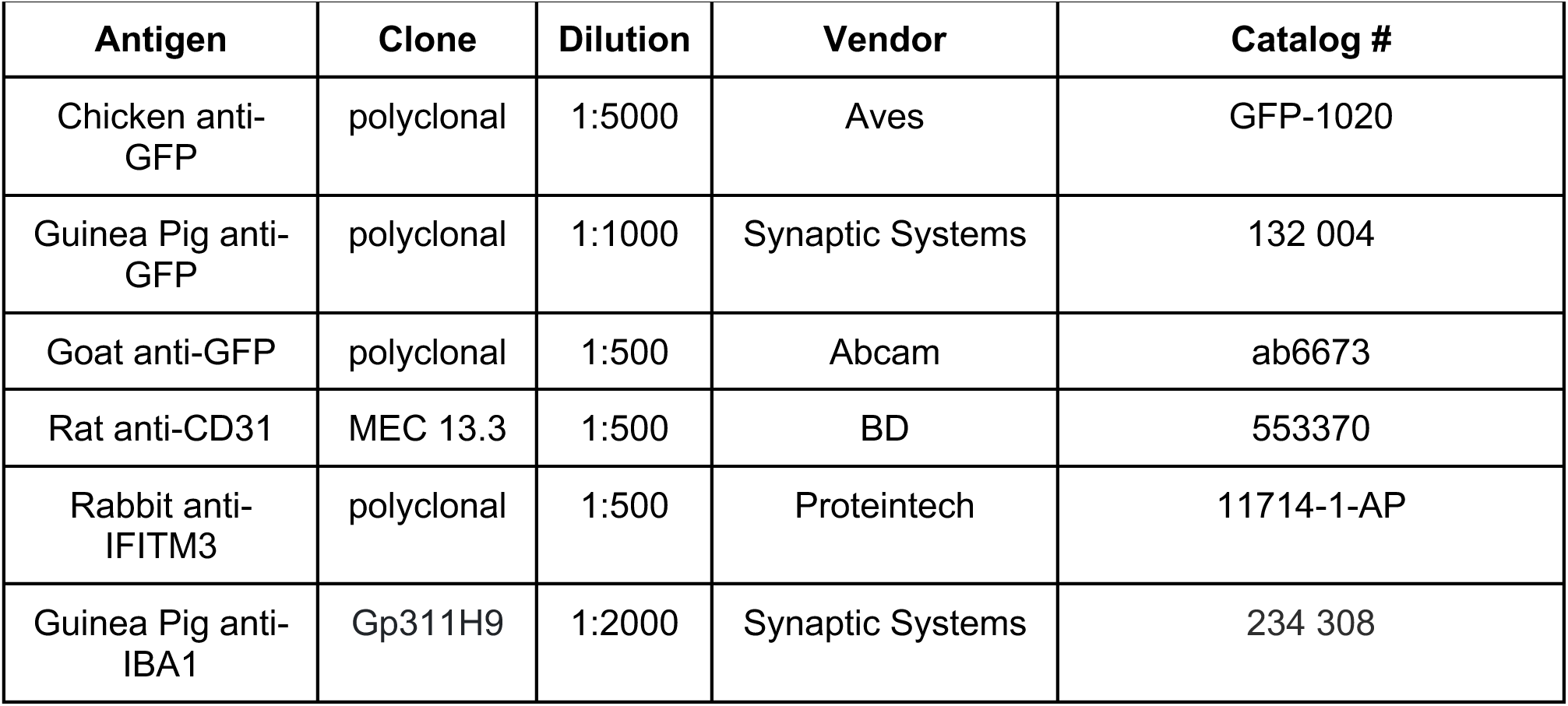
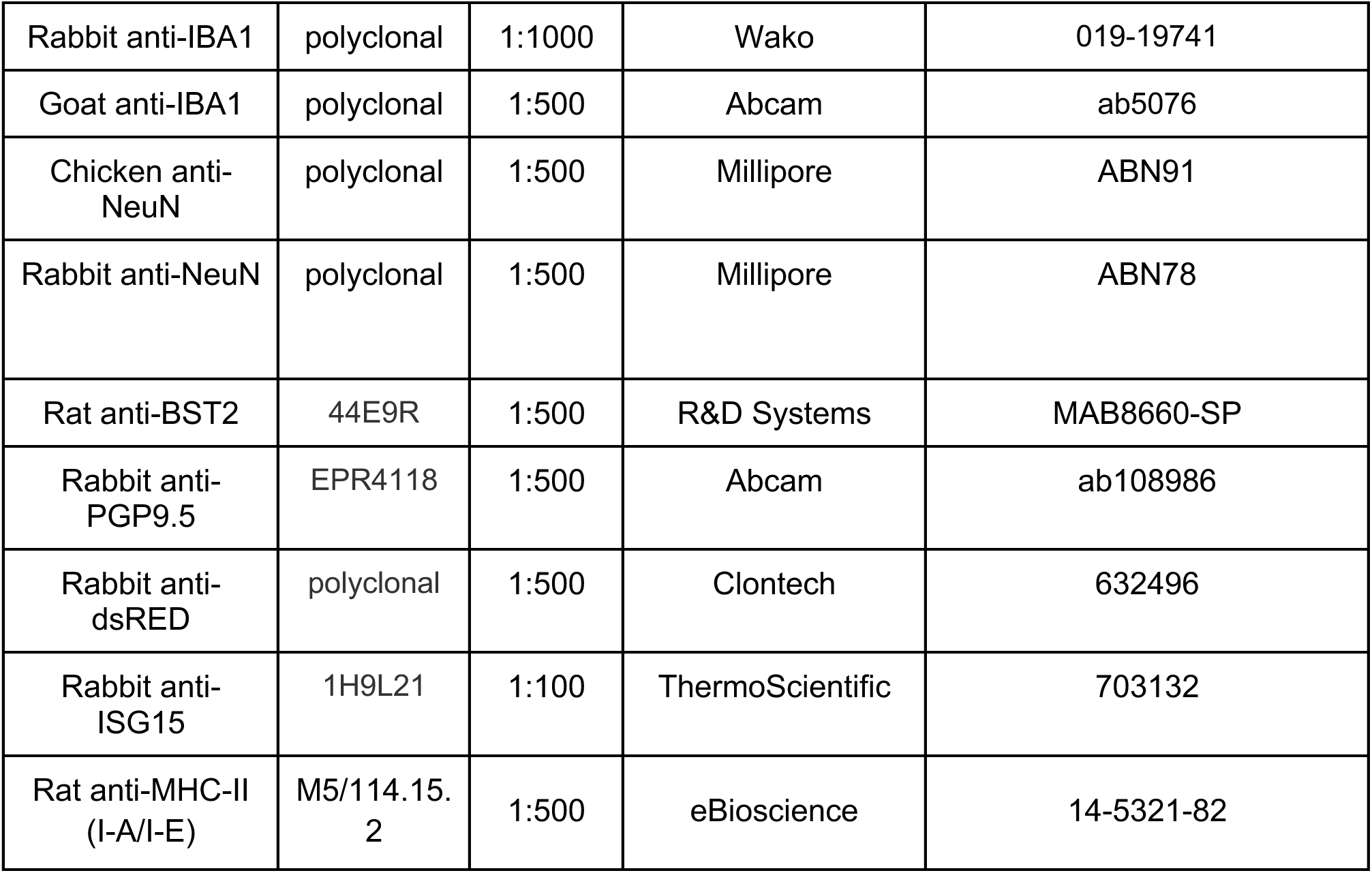

*Antibodies for immunofluorescence:*
| Antigen | Clone | Dilution | Vendor | Catalog # |
| --- | --- | --- | --- | --- |
| Chicken anti-GFP | polyclonal | 1:5000 | Aves | GFP-1020 |
| Guinea Pig anti-GFP | polyclonal | 1:1000 | Synaptic Systems | 132 004 |
| Goat anti-GFP | polyclonal | 1:500 | Abcam | ab6673 |
| Rat anti-CD31 | MEC 13.3 | 1:500 | BD | 553370 |
| Rabbit anti-IFITM3 | polyclonal | 1:500 | Proteintech | 11714-1-AP |
| Guinea Pig anti-IBA1 | Gp311H9 | 1:2000 | Synaptic Systems | 234 308 |
| Rabbit anti-IBA1 | polyclonal | 1:1000 | Wako | 019-19741 |
| Goat anti-IBA1 | polyclonal | 1:500 | Abcam | ab5076 |
| Chicken anti-NeuN | polyclonal | 1:500 | Millipore | ABN91 |
| Rabbit anti-NeuN | polyclonal | 1:500 | Millipore | ABN78 |
| Rat anti-BST2 | 44E9R | 1:500 | R&D Systems | MAB8660-SP |
| Rabbit anti-PGP9.5 | EPR4118 | 1:500 | Abcam | ab108986 |
| Rabbit anti-dsRED | polyclonal | 1:500 | Clontech | 632496 |
| Rabbit anti-ISG15 | 1H9L21 | 1:100 | ThermoScientific | 703132 |
| Rat anti-MHC-II (I-A/I-E) | M5/114.15.2 | 1:500 | eBioscience | 14-5321-82 |

